# An essential and highly selective protein import pathway encoded by nucleus-forming phage

**DOI:** 10.1101/2024.03.21.585822

**Authors:** Chase J. Morgan, Eray Enustun, Emily G. Armbruster, Erica A. Birkholz, Amy Prichard, Taylor Forman, Ann Aindow, Wichanan Wannasrichan, Sela Peters, Koe Inlow, Isabelle L. Shepherd, Alma Razavilar, Vorrapon Chaikeeratisak, Benjamin A. Adler, Brady F. Cress, Jennifer A. Doudna, Kit Pogliano, Elizabeth Villa, Kevin D. Corbett, Joe Pogliano

**Author notes:** Corresponding author 4111 Natural Sciences Building University of California San Diego La Jolla, CA 92093, USA. **Competing interests:**. K.P. and J.P. have an equity interest in Linnaeus Bioscience Incorporated and receive income. The terms of this arrangement have been reviewed and approved by the University of California, San Diego, in accordance with its conflict-of-interest policies. The Regents of the University of California have patents issued and pending for CRISPR technologies on which J.A.D., B.F.C., B.A.A., J.P., K.P., and E.G.A. are inventors. J.A.D. is a cofounder of Caribou Biosciences, Editas Medicine, Scribe Therapeutics, Intellia Therapeutics, and Mammoth Biosciences. J.A.D. is a scientific advisory board member of Vertex, Caribou Biosciences, Intellia Therapeutics, Scribe Therapeutics, Mammoth Biosciences, Algen Biotechnologies, Felix Biosciences, The Column Group, and Inari. J.A.D. is Chief Science Advisor to Sixth Street, a Director at Johnson & Johnson, Altos and Tempus, and has research projects sponsored by Apple Tree Partners and Roche.

## Abstract

Targeting proteins to specific subcellular destinations is essential in prokaryotes, eukaryotes, and the viruses that infect them. Chimalliviridae phages encapsulate their genomes in a nucleus-like replication compartment composed of the protein chimallin (ChmA) that excludes ribosomes and decouples transcription from translation. These phages selectively partition proteins between the phage nucleus and the bacterial cytoplasm. Currently, the genes and signals that govern selective protein import into the phage nucleus are unknown. Here we identify two components of this novel protein import pathway: a species-specific surface-exposed region of a phage intranuclear protein required for nuclear entry and a conserved protein, PicA, that facilitates cargo protein trafficking across the phage nuclear shell. We also identify a defective cargo protein that is targeted to PicA on the nuclear periphery but fails to enter the nucleus, providing insight into the mechanism of nuclear protein trafficking. Using CRISPRi-ART protein expression knockdown of PicA, we show that PicA is essential early in the chimallivirus replication cycle. Together our results allow us to propose a multistep model for the Protein Import Chimallivirus (PIC) pathway, where proteins are targeted to PicA by amino acids on their surface, and then licensed by PicA for nuclear entry. The divergence in the selectivity of this pathway between closely-related chimalliviruses implicates its role as a key player in the evolutionary arms race between competing phages and their hosts.

**Significance Statement:** The phage nucleus is an enclosed replication compartment built by Chimalliviridae phages that, similar to the eukaryotic nucleus, separates transcription from translation and selectively imports certain proteins. This allows the phage to concentrate proteins required for DNA replication and transcription while excluding DNA-targeting host defense proteins. However, the mechanism of selective trafficking into the phage nucleus is currently unknown. Here we determine the region of a phage nuclear protein that targets it for nuclear import and identify a conserved, essential nuclear shell-associated protein that plays a key role in this process. This work provides the first mechanistic model of selective import into the phage nucleus.

## Introduction

Compartmentalization of the cytoplasm into organelles enclosed by a semipermeable barrier is a hallmark of eukaryotic life. The phage nucleus is a viral organelle produced by phages in the Chimalliviridae family and the first DNA-enclosing viral replication compartment discovered in a prokaryotic cell (1–4). During the chimallivirus infection cycle, the phage genomes replicate within the phage nucleus, which is bounded by a lattice composed mainly of the phage protein chimallin (ChmA) (1–5). The ChmA barrier excludes ribosomes, decoupling transcription from translation, so macromolecular transport across the nuclear shell is necessary (1, 5). mRNA must be exported, while many proteins for DNA replication and transcription must be imported. The ChmA lattice is porous to metabolites and nucleotides required for transcription, but these pores are too small to accommodate trafficking of folded macromolecules (5, 6). While a putative nuclear shell mRNA transporter was recently discovered (7), no protein transporter has been identified.

Protein import across the ChmA barrier is selective (1–4, 8–10). Certain phage proteins concentrate in the nucleus including RNA polymerase subunits, a RecA-related recombinase (henceforth RecA), DNA polymerase, and a number of proteins of unknown function (1, 3, 4, 11, 12). In contrast, many phage proteins are excluded and remain in the host cytoplasm including metabolic enzymes like thymidylate kinase and virion structural proteins (1, 3, 4, 11). Many host proteins are also excluded, including DNA-targeting defense proteins such as Cas9 and restriction endonucleases (1, 4, 8, 9), which provides the phage with broad protection against many host defenses.

Furthermore, during co-infection of a single cell by two different nucleus-forming phages, the phage nucleus of one species excludes a toxic nuclease produced by the other (13, 33). The high degree of specificity seen in protein localization to the phage nucleus has led to the hypothesis that chimalliviruses encode a selective protein trafficking system (1, 2, 4, 10, 13, 14).

In eukaryotes, proteins can be sorted into organelles including the endoplasmic reticulum, nucleus, mitochondria, and peroxisomes (15–20). In prokaryotes, proteins can be translocated to the periplasm, outer membrane, or secreted outside of the cell (21–28). While the mechanisms underlying these transport systems vary significantly, most rely on a signal sequence encoded within the cargo protein, a means of recognizing that signal, and a protein complex to facilitate the trafficking of the cargo to its destination (21–28). In nucleus-forming phages, it is reasonable to expect that protein trafficking would follow a similar pattern. However, bioinformatic analyses of nuclear proteins have failed to identify any conserved signal sequences.

Experiments with GFPmut1 suggested that residues on the protein’s surface are essential for its nuclear import (10). GFPmut1 is a GFP variant that is imported into the PhiKZ phage nucleus but not the nuclei of other phages, while other GFP variants tested are not imported into any phage nucleus. Import of GFPmut1 is lost if a surface-exposed phenylalanine (F99) is mutated and partially lost if a nearby surface-exposed methionine (M153) is mutated (10), suggesting the signals that target proteins to the nucleus may be encoded in amino acids distant in sequence but located on the same surface in the folded protein. However, the signals needed for targeting endogenous phage proteins to the nucleus remain unidentified.

Here we use genetic and cell biological approaches to determine the recognition region on a phage protein that targets it for nuclear import and demonstrate that a conserved protein in nucleus-forming phages, which we term PicA (**P**rotein **i**mporter of **c**himalliviruses **A**), plays a key role in selective nuclear import. PicA is the first protein identified to be involved in selective protein trafficking into the phage nucleus, and we show that it is essential for viral reproduction. These results provide insight into how these phages selectively partition proteins between the phage nucleus and the bacterial cytoplasm, and establish the PIC pathway as an essential component of the phage replication cycle with direct implications for phage-phage and phage-host competition.

## Results

### Protein localization to the phage nucleus is species- and sequence-specific

To better understand the specificity of the phage nuclear import machinery, we studied the ability of PhiPA3 and PhiKZ, two closely related *Pseudomonas* phages, to import each other’s proteins. We expressed five previously identified PhiPA3 nuclear proteins (11) with a C-terminal sfGFP tag (RecA, gp108, gp200, gp78, and gp257) in *Pseudomonas aeruginosa* and infected the cells with either PhiPA3 or PhiKZ (sfGFP alone does not localize to the phage nucleus of either phage (10)). As expected, all five PhiPA3 proteins localized within the PhiPA3 nucleus (Fig 1A). When cells were infected with PhiKZ, PhiPA3 RecA was imported into the PhiKZ nucleus, while the other four PhiPA3 proteins were excluded from the PhiKZ nucleus (Fig 1A), demonstrating that PhiKZ does not import the majority of PhiPA3 nuclear proteins.

**Figure 1.**
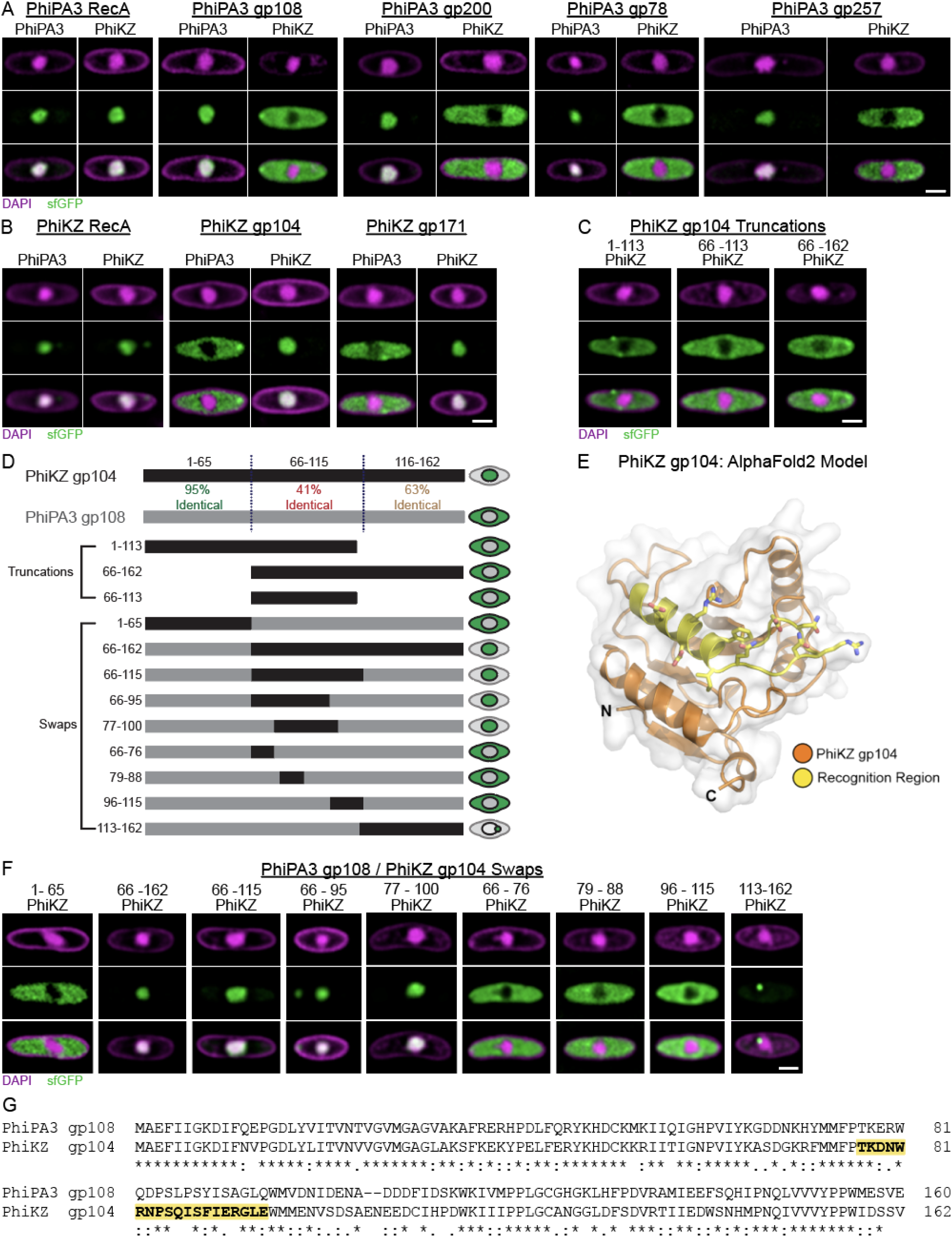
Protein trafficking into the phage nucleus is species- and sequence-specific. A) *P. aeruginosa* strain PA01 K2733 cells expressing the indicated sfGFP-tagged PhiPA3 nuclear protein infected with either PhiPA3 or PhiKZ. DAPI staining (purple) shows the phage nucleus in the center of the cell and peptidoglycan at the cell periphery. sfGFP (green) shows the localization of the tagged protein either to the phage nucleus or the cytoplasm. B) *P. aeruginosa* cells expressing the indicated sfGFP-tagged PhiKZ nuclear protein infected with either PhiPA3 or PhiKZ. C) *P. aeruginosa* cells expressing the indicated sfGFP-tagged PhiKZ gp104 truncation infected with PhiKZ D) Schematic depicting the percent conservation between PhiKZ gp104 and PhiPA3 gp108 and the sfGFP-tagged protein constructs expressed in (D) and (E) with their phenotype indicated on the right. Numbers on the left indicate the amino acids belonging to PhiKZ gp104 in each construct. E) AlphaFold predicted structure of PhiKZ gp104 with the experimentally determined recognition region required for nuclear localization shown in yellow F) *P. aeruginosa* cells expressing the indicated sfGFP-tagged PhiKZ gp104 / PhiPA3 gp108 chimera infected with PhiKZ. All infections were imaged at 30-45 minutes post-infection. All scale bars are 1 µm. G) Clustal Omega generated protein sequence alignment of PhiPA3 gp108 and PhiKZ gp104. “*” denotes conserved amino acids. “:” and “.” denotes highly similar and weakly similar amino acids respectively. Sequence of PhiKZ gp104 import recognition region is highlighted in yellow and bolded.

Three of the five PhiPA3 proteins have a homolog in PhiKZ: PhiKZ RecA, gp104, and gp171. As expected, these three PhiKZ proteins localize to the PhiKZ nucleus (Fig 1B). PhiKZ RecA was also imported into the PhiPA3 nucleus. However, PhiKZ gp104 and gp171 were excluded from the PhiPA3 nucleus (Fig 1B), demonstrating that the PhiPA3 nucleus excludes the majority of PhiKZ nuclear proteins as well. These data show that these phages primarily import their cognate proteins and exclude the proteins of other phages from their nucleus.

PhiKZ and PhiPA3 RecA share the highest sequence identity of the proteins tested (∼80% amino acid sequence identity) (Fig S1D), but the other homolog pairs also share considerable amino acid sequence identity. PhiKZ gp104 and PhiPA3 gp108 share ∼65%, while PhiKZ gp171 and PhiPA3 gp200 share ∼54% (Fig 1F, Fig S1C). The fact that these latter proteins only localize to their cognate nucleus demonstrates that these phages can discriminate between closely related homologs with significant sequence identity, suggesting the existence of phage-specific import recognition signals.

We investigated this hypothesis with PhiKZ gp104, a 162-amino acid protein of unknown function belonging to the conserved macro-domain family (29). We first expressed truncated versions of sfGFP-tagged PhiKZ gp104 in *P. aeruginosa* and infected with PhiKZ (Fig 1C and D). While full length PhiKZ gp104 was imported (Fig 1B), none of the truncations localized to the phage nucleus, suggesting that multiple distant sequences are required for import or that the phage recognizes the surface of a properly folded protein.

Next, we generated chimeras of PhiKZ gp104 and its homolog PhiPA3 gp108 and tested for localization to the PhiKZ nucleus to identify a region conferring import specificity (Figs 1D and F). PhiKZ gp104 and PhiPA3 gp108 have nearly identical N-termini (amino acids 1-65) and are most divergent in the middle and C-terminal regions (Figs 1D and G). Since PhiPA3 gp108 does not localize to the PhiKZ nucleus (Fig 1A) but shares high sequence and structural homology to PhiKZ gp104 (Fig 1G, Fig S1A), this approach allowed us to screen for sequence-specific localization effects while still maintaining the full sequence length and general structure of the protein. As expected, fusing the N-terminal 65 amino acids of PhiKZ gp104 to the C-terminal 95 amino acids of PhiPA3 gp108 maintained the cytoplasmic localization of the chimeric PhiPA3 gp108 during PhiKZ infection (Fig 1F). However, swapping the C-terminal ∼2/3rds (amino acids 66-162) or the middle ∼1/3 (amino acids 66-115) of PhiKZ gp104 into the homologous region of PhiPA3 gp108 caused the chimera to localize to the nucleus during PhiKZ infection (Fig 1F). Shortening the region of PhiKZ gp104 swapped into PhiPA3 gp108 revealed two more chimeras that localize to the PhiKZ nucleus: chimeras containing PhiKZ gp104 amino acids 66-95 and PhiKZ gp104 amino acids 77-100 (Fig 1F). In comparison, three chimeras with shorter sequences from PhiKZ gp104 did not localize to the PhiKZ nucleus (Fig 1F). These data indicate that at least part of the region between amino acids 77 and 95 is essential for PhiKZ gp104 localization to the phage nucleus, but the region between amino acids 79-88 is either insufficiently short or unnecessary. The region between amino acids 77 and 95 is ∼47% identical between the homologs, demonstrating that the phage import machinery recognizes only a small number (≤10) of non-conserved amino acids (Fig S1B).

Mapping the non-conserved amino acids in this region to the AlphaFold-predicted structure (30) of PhiKZ gp104 shows that they are part of an exposed surface of the protein that we call the recognition region (Fig 1E). Aligning the sequence of this region with other known PhiKZ nuclear proteins including RecA, RNA polymerases, and gp171 did not reveal a conserved motif (31). These results indicate that protein trafficking into the phage nucleus likely involves a recognition surface rather than an N- or C-terminal signal sequence as found in other import systems (15, 17, 19, 22, 24). This is consistent with previous results that two surface-exposed amino acids which are distant in the primary sequence are essential for the localization of GFPmut1 to the PhiKZ nucleus (10).

One sfGFP-tagged chimera containing the first 112 amino acids of PhiPA3 gp108 with the C-terminal segment of PhiKZ gp104 (amino acids 113-162) produced an unexpected phenotype. During PhiKZ infection, this protein accumulates in a single punctum at the periphery of the phage nucleus but fails to enter (Fig 1F). This suggests there are two major steps in the import process, nuclear targeting and translocation across the nuclear shell, and that the chimera likely represents a stalled transport intermediate. The presence of a single focus instead of multiple foci spread around the nuclear shell additionally suggests that the nucleus has a single site of protein import.

### Mutations in the conserved shell-associated protein PicA alter selective protein trafficking into the phage nucleus

To determine the identity of the factors that facilitate protein trafficking into the phage nucleus, we designed a genetic selection to isolate mutant phage that no longer import a toxic protein (32). We expressed a fusion between GFPmut1 and the PhiPA3 endonuclease gp210 and then infected with PhiKZ (Fig 2A). GFPmut1 can be used to drive PhiKZ nuclear import of proteins that are normally excluded and remain in the host cytoplasm (10, 13, 33). PhiPA3 gp210 is normally excluded from the PhiKZ phage nucleus, but gp210-GFPmut1 is imported, where it becomes toxic by cleaving the PhiKZ genome and preventing replication (13, 33). We therefore predicted that through selection for phage that can replicate in the presence of gp210-GFPmut1, we could isolate escape mutants that have altered the ability to traffic proteins into the phage nucleus (Fig 2A).

**Figure 2.**
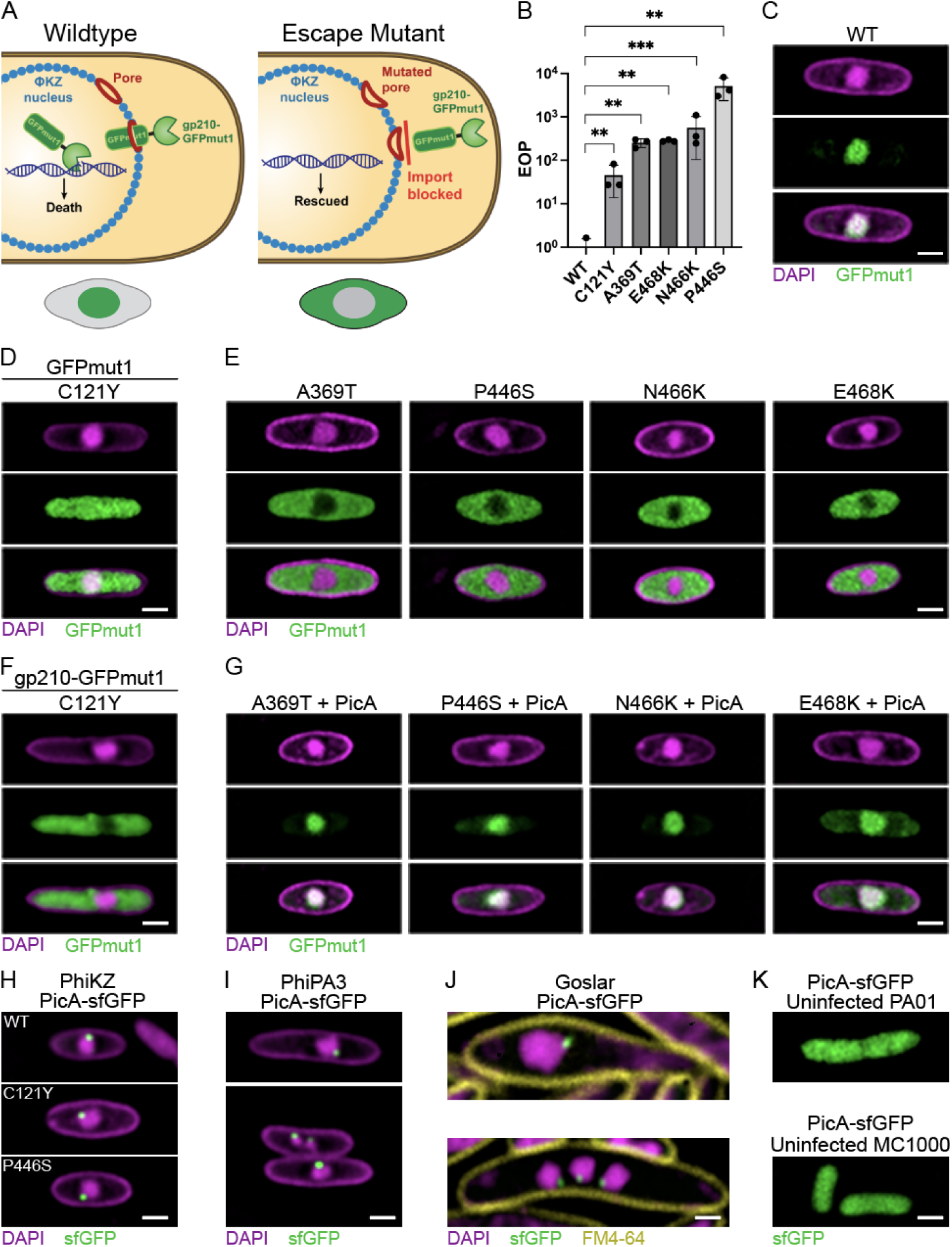
The nuclear-shell associated protein PicA plays a key role in selective protein trafficking into the phage nucleus. A) Schematic illustrating the genetic selection performed with the toxic fusion protein gp210-GFPmut1 targeted to the PhiKZ nucleus with the expected GFPmut1 localization phenotype of the wildtype and mutant PhiKZ depicted below. B) Mean efficiency of plating of mutant PhiKZ relative to wildtype calculated by spot titer on lawns of *P. aeruginosa* expressing gp210-GFPmut1. Error bars indicate standard deviations. ** = p<0.01, *** = p<0.001. Significance determined by paired T-test. C) Microscopy image of *P. aeruginosa* expressing GFPmut1 infected with wildtype PhiKZ. All *P. aeruginosa* phage infections imaged at 30-45 minutes post-infection. D) *P. aeruginosa* expressing GFPmut1 infected with PicA C121Y mutant PhiKZ. E) *P. aeruginosa* expressing GFPmut1 infected with PhiKZ containing the indicated mutation in PicA. F) *P. aeruginosa* expressing gp210-GFPmut1 infected with PicA C121Y mutant PhiKZ. G) *P. aeruginosa* expressing both wildtype PicA and GFPmut1 infected with the indicated PicA mutant PhiKZ. H) *P. aeruginosa* expressing sfGFP-tagged wildtype or mutant PicA infected with wildtype PhiKZ I) *P. aeruginosa* expressing sfGFP-tagged wildtype PicA from phage PhiPA3 infected with PhiPA3. Upper panel shows a cell with a single nucleus, while the lower panel depicts a cell with multiple phage nuclei and a single PicA punctum on each nucleus. J) *E. coli* strain MC1000 expressing sfGFP-tagged Goslar wildtype PicA infected with Goslar. Upper panel shows a cell with a single nucleus, while the lower panel depicts a cell with multiple phage nuclei and a single PicA punctum on each nucleus. Images taken at 75-90 minutes post infection. K) PhiKZ PicA-sfGFP expressed in an uninfected *P. aeruginosa* PA01 K2733 cell (left) or Goslar PicA-sfGFP expressed in an uninfected *E. coli* MC1000 cell (right). All scale bars are 1 µm.

Strains expressing gp210-GFPmut1 exhibit a ∼10,000-fold reduction in PhiKZ plaquing, and mutants that escape this selection can be isolated (33). PhiKZ mutants that escape gp210-GFPmut1 selection have mutations in one of two genes with approximately equal frequency (Table 1). The first is the virion RNA polymerase subunit gp178 gene, which contains the previously identified gp210 nuclease target site (33). Mutations at this site are expected to reduce or eliminate gp210-mediated cleavage, rendering the mutant resistant to gp210. The second mutated gene encodes a protein of unknown function, gp69. Five independent mutant phages with a single mutation in gp69 were isolated and each of these mutants showed impaired protein localization to the phage nucleus, leading us to name this protein PicA (**P**rotein **i**mporter of **c**himalliviruses **A**) (Figs 2D, E, and F). These mutants show a ∼50 to ∼5,000 fold rescue in titer compared to wildtype when plaquing on cells expressing gp210-GFPmut1 (Fig 2B). Live cell imaging of infected cells showed that four of the five isolated mutants (A369T, P446S, N466K, and E468K) exclude GFPmut1 from the PhiKZ nucleus (Fig 2E). The fifth mutant (C121Y) does not fully exclude GFPmut1 from the nucleus (Fig 2D), but does exclude the gp210-GFPmut1 fusion protein from the nucleus (Fig 2F). Infections with all of these mutant phages appear to progress normally and still concentrate sfGFP-tagged PhiKZ nuclear proteins (Fig S2), suggesting that import of endogenous phage proteins is not strongly affected by these point mutations. Furthermore, expressing wildtype PicA in conjunction with GFPmut1 and infecting with mutant phage restores the nuclear localization of GFPmut1, indicating that the effect is specific to the mutations in PicA and that these mutations are recessive to the wildtype protein (Fig 2G). We tested to see if we could perform a similar selection with a PhiKZ nuclear protein, however selecting mutants with PhiKZ gp104 fused to PhiPA3 gp210 only produced escape mutants that had altered the nuclease target site in gp178 (Table 1). This suggests that complete loss of nuclear import of an endogenous phage protein, gp104, was more detrimental to phage fitness than mutating the nuclease target site.

**Table 1.** PhiKZ mutants isolated in this study. “n” indicates any mutation from the wildtype base. X indicates any amino acid substitution from the wildtype residue. Phenotypes related to nuclear protein trafficking are bolded.

| Method of Isolation | Gene Product Mutated | DNA Mutation | Amino Acid Mutation | Frequency | Phenotype |
| --- | --- | --- | --- | --- | --- |
| gp210-GFPmut1 Selection | gp178 | a3215n | D1072X | 18/32 | Resistant to Nuclease Activity |
|  | PicA (gp69) | g362a | C121Y | 1/32 | **Excludes gp210-GFPmut1 from Phage Nucleus, but not GFPmut1** |
|  | PicA (gp69) | g1105a | A369T | 1/32 | **Excludes GFPmut1 from Phage Nucleus** |
|  | PicA (gp69) | c1336t | P446S | 8/32 | **Excludes GFPmut1 from Phage Nucleus** |
|  | PicA (gp69) | t1398g | N466K | 3/32 | **Excludes GFPmut1 from Phage Nucleus** |
|  | PicA (gp69) | g1402a | E468K | 1/32 | **Excludes GFPmut1 from Phage Nucleus** |
| gp210-gp104 Selection | gp178 | a3215n | D1072X | 9/10 | Resistant to Nuclease Activity |
|  | gp178 | a3212g | D1071G | 1/10 | Resistant to Nuclease Activity |
| Cas13a PicA g3 | PicA (gp69) | t222g | R74R | 1/7 | No Phenotype Observed |
|  | PicA (gp69) | g221a | R74H | 5/7 | No Phenotype Observed |
|  | PicA (gp69) | c224g | A75G | 1/7 | No Phenotype Observed |
| Cas13a PicA g4 | PicA (gp69) | t1077a | H359Q | 4/17 | No Phenotype Observed |
|  | PicA (gp69) | g1081c | G361C | 3/17 | No Phenotype Observed |
|  | PicA (gp69) | t1092g | D364E | 1/17 | No Phenotype Observed |
|  | PicA (gp69) | t1095c | H365H | 5/17 | No Phenotype Observed |
|  | PicA (gp69) | t1095a | H365Q | 1/17 | **Excludes GFPmut1 from Phage Nucleus** |
|  | PicA (gp69) | a1097c | H366P | 2/17 | No Phenotype Observed |
|  | PicA (gp69) | a1099c | T367P | 1/17 | **Delayed Nuclear Localization** |
| Cas13a PicA g6 | PicA (gp69) | c889a | Q297K | 10/10 | **Altered gp104 Nuclear Localization** |

The *picA* gene is part of the Chimalliviridae core genome and is conserved in all known phages that encode the nuclear shell protein ChmA (4) (Fig S3). Furthermore, the PicA homolog from phage PhiPA3 (gp63) was recently identified as nuclear shell-associated and part of the ChmA interaction network (11). Live cell imaging of infected cells expressing sfGFP-tagged PicA shows that PicA from diverse phages, including PhiKZ, PhiPA3, and *E. coli* phage Goslar (3, 34), localizes to the nuclear periphery (Figs 2H-J, Fig S4). For example, PhiKZ PicA-sfGFP localizes into a single punctum at the nuclear periphery during PhiKZ infection (Fig 2I) but is diffuse in uninfected cells (Figs 2H and K). This localization is not affected by mutations (C121Y and P446S) in PicA that alter GFPmut1 import (Fig 2H). PhiPA3 PicA (Fig 2I) and Goslar PicA (gp174) (Fig 2J) also localize in a single punctum at the nuclear periphery during infection of their respective hosts, *P. aeruginosa* and *E. coli*. When infected cells display more than one nucleus during infection, a single PicA-sfGFP punctum can be seen at the periphery of each nucleus (Figs 2I and J), suggesting that this phenotype is biologically relevant and not due to non-specific aggregation of the sfGFP-tagged protein. Together, these results suggest that PicA is a conserved protein that interacts with the nuclear shell for the purpose of trafficking proteins across the ChmA lattice.

To gain deeper understanding of the key structural or sequence elements required for PicA to modulate protein import, we generated a series of *picA* mutants with LbuCas13a (*Leptotrichia buccalis* Cas13a, henceforth Cas13a) using guides targeting specific regions of PhiKZ PicA (35). Cas13a uses a guide RNA to recognize specific phage encoded RNAs, which activates its nonspecific RNA nucleolytic activity. This leads to degradation of both phage and host cellular RNA, cell dormancy, and phage restriction (36) (Fig 3A). Cas13a produces a strong selection against the wildtype gene so only phage with mutations in the targeted region can replicate (35, 37). RNA-targeting Cas systems are particularly well-suited for counterselection of chimalliviruses, as their DNA is protected within the phage nucleus, but their mRNA is not (8, 9, 35, 37). Regions of the *picA* gene to be targeted with guides were chosen based upon the AlphaFold2 prediction of the PicA structure (Fig S7). Sequences proximal to the previously generated mutants, as well as sequences corresponding to additional faces of the protein were targeted with guide RNAs containing 31-nucleotide spacer sequences. We transformed plasmids expressing both Cas13a and single guides against *picA* into *P. aeruginosa* and infected with wildtype PhiKZ. In doing so, we selected for spontaneous phage mutants that escape guide recognition and subsequently screened them for protein import defects. Of the seven guides tested, three (guide 3, 4, and 6) produced isolable escape mutants. Among the 34 PhiKZ mutants we isolated, we observed 11 unique mutations in the *picA* gene (Table 1). Two were silent mutations, while the other nine resulted in single amino acid substitutions. Six of these nine mutants showed no change in their ability to import nuclear proteins. One of the remaining three mutants (H365Q) showed a defect only in GFPmut1 localization (Fig 3C), demonstrating that we can recapitulate the previously identified phenotype (Fig 2E) with this selection method.

**Figure 3.**
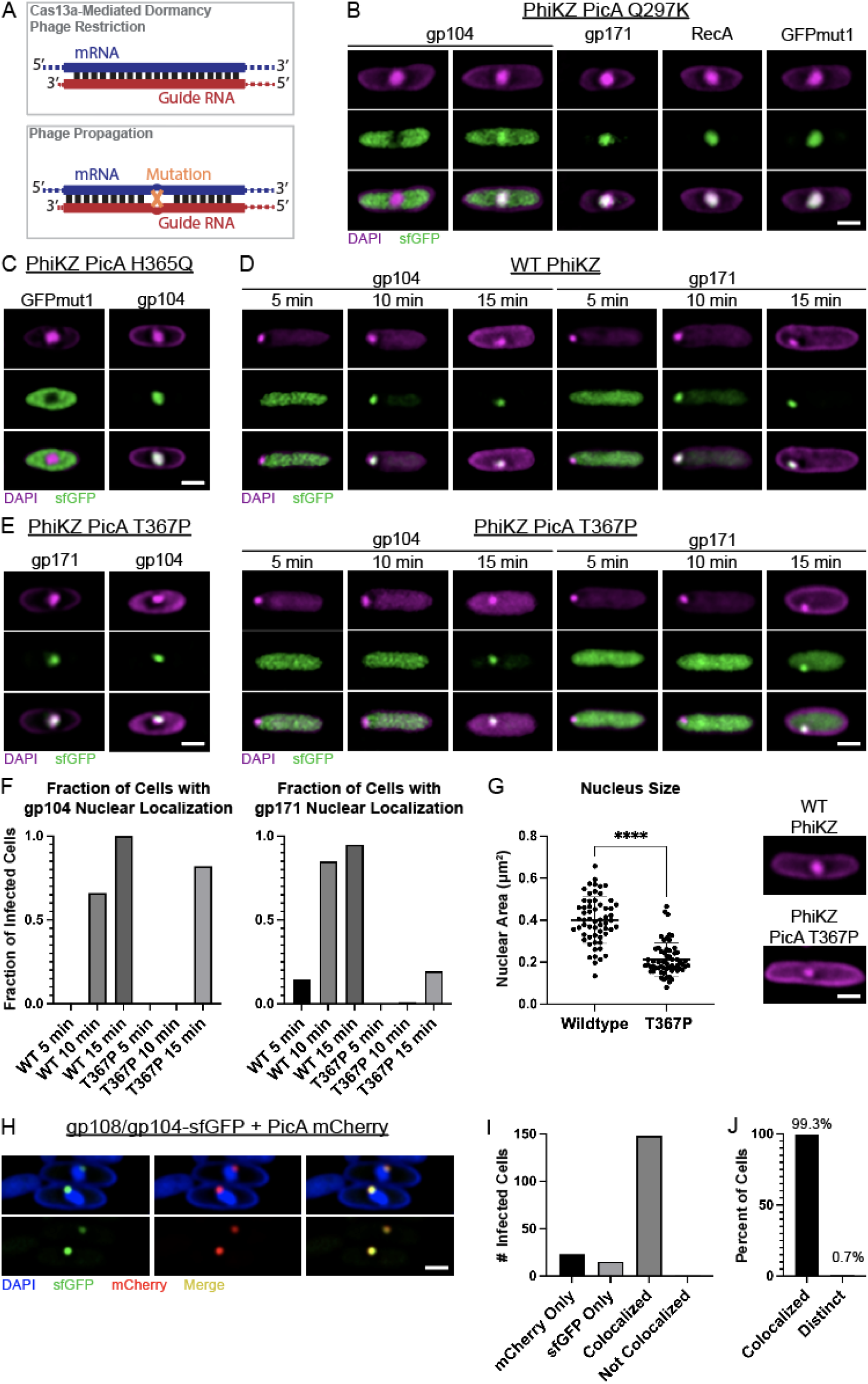
PicA traffics phage proteins into the phage nucleus. A) Schematic of the mechanism of mutant isolation with catalytically-active Cas13a. Anti-sense guide RNAs pair with mRNA of expressed genes and activate Cas13a, which leads to cell dormancy and phage restriction. Mutant phages that exist in the population escape targeting and become the dominant phage in the population. B) *P. aeruginosa* cells expressing the indicated sfGFP-tagged PhiKZ protein infected with PhiKZ PicA Q297K mutant. Images taken 30-45 minutes post infection. C) *P. aeruginosa* cells expressing GFPmut1 (left) or sfGFP-tagged PhiKZ gp104 (right) infected with PhiKZ PicA H365Q mutant imaged at 30 minutes post infection. D) *P. aeruginosa* cells expressing sfGFP-tagged PhiKZ gp104 (left) or sfGFP-tagged gp171 (right) infected with PhiKZ wildtype or PicA T367P mutant. Images taken after 5 minutes, 10 minutes, or 15 minutes of incubation with phage. E) *P. aeruginosa* cells infected with wildtype or PicA T367P mutant PhiKZ imaged at 30 minutes post infection. F) Quantification of the fraction of infected cells that displayed a nuclear localization of the specified protein at the indicated time point as shown in (D). Nuclear localization during infection was called if the maximum GFP signal co-localized with the DAPI-stained phage genome. Localization was called for the total number of infected cells imaged for each condition (n= 68-159 per condition). G) DAPI-stained wildtype *P. aeruginosa* cells infected with wildtype or PicA T367P mutant PhiKZ at 30 minutes post infection with quantification of nuclear area for cells infected with wildtype or PicA T367P mutant PhiKZ imaged. Line indicates mean and error bars indicate standard deviation. N=60 per condition. **** = p<0.0001 by Mann-Whitney test. All scale bars are 1 µm. H) *P. aeruginosa* cells co-expressing a stalled PhiKZ gp104/ PhiPA3 gp108 chimera (PhiPA3gp108[PhiKZgp104(113–162)]) with sfGFP and PicA tagged with mCherry infected with wildtype PhiKZ. DAPI stain is shown in blue, GFP in green, and mCherry in red. Co-localization of mCherry and sfGFP can be seen in yellow. Lower panels depict only the fluorescent protein. I) Number of infected cells co-expressing both sfGFP-tagged chimera and PicA-mCherry that have fluorescence in at least one channel above threshold displaying the indicated phenotype. N = 187 cells (mCherry only = 23, sfGFP only = 15, mCherry/sfGFP with colocalization = 148, mCherry/sfGFP without colocalization = 1) J) Percent of infected cells with both mCherry and sfGFP fluorescence above threshold that display either overlapping (colocalized, 99.3%) or non-overlapping (distinct, 0.7%) fluorescent puncta at the nuclear periphery. N = 149 cells. See Fig S5 for more detail.

The remaining two mutants (Q297K and T367P) showed a defect in the nuclear localization of one or more sfGFP-tagged phage proteins, implicating PicA in the endogenous protein trafficking pathway. PicA Q297K mutant PhiKZ fails to fully concentrate sfGFP-tagged PhiKZ gp104 in the phage nucleus after 30 minutes post-infection (mpi), while nuclear localization of PhiKZ gp171, PhiKZ RecA, or GFPmut1 are unaffected (Fig 3B). PicA T367P mutant PhiKZ ultimately concentrates both PhiKZ gp104 and PhiKZ gp171 in the nucleus but shows a temporal delay in protein localization and produces smaller nuclei (Figs 3D-G). At 5 mpi, the phage genome can be seen as a DAPI-stained punctum at the pole of the cell in both wildtype and PicA T367P mutant PhiKZ infections. At this time point, exogenously expressed sfGFP-tagged PhiKZ gp104 and gp171 are diffuse in the cytoplasm (Figs 3D and F). By 10 mpi, sfGFP-tagged PhiKZ gp104 and gp171 are predominantly localized to the wildtype phage nucleus, while the fluorescence signal from both proteins is still diffuse in the cytoplasm during infection by the mutant. PhiKZ gp104-sfGFP and gp171-sfGFP do not concentrate in the mutant nucleus until 15 mpi or later (Figs 3D-F), although both proteins are predominantly localized within the phage nucleus by 30 mpi (Fig 3E). However, at 30 mpi PicA T367P mutant nuclei are significantly smaller than wildtype nuclei, indicating a delay in nuclear development and that protein import is required for nucleus growth (Fig 3G). Together, these two mutations (Q297K and T367P) implicate PhiKZ PicA in the trafficking of endogenous phage proteins into the phage nucleus.

### The PicA import machinery colocalizes with a transport intermediate

Since mutations in PicA alter selective protein trafficking, we sought to determine if PicA associates with imported proteins during phage infection. One sfGFP-tagged chimera of PhiPA3 gp108 and PhiKZ gp104 (PhiPA3gp108-[PhiKZ gp104(113–162)]-sfGFP) is diffuse in the cytoplasm of uninfected cells (Fig S6) but accumulates in a single punctum on the phage nuclear shell during infection, likely representing a stalled transport intermediate (Fig 1F). Notably, fluorescently-tagged PicA also localizes in a single punctum on the phage nucleus surface (Fig. 2I-J). If PicA is part of the import machinery, we would expect it to colocalize with the putative transport intermediate. Therefore, we infected cells co-expressing the sfGFP-tagged stalled chimera and mCherry-tagged PicA with PhiKZ (Fig 3H). Of the 187 infected cells observed, 149 contained puncta of both PicA-mCherry and stalled chimera-sfGFP, while the other 38 only contained a detectable punctum of one fluorescently tagged protein (Fig 3I). Strikingly, in cells where both fluorescent fusions were detected, PicA-mCherry and stalled chimera-sfGFP colocalize on the nuclear periphery in 99.3% of infections (n = 149). (Figs 3H and 3J, Fig S5). This provides evidence that PicA forms part of the import machinery through which imported proteins pass and supports a mechanism in which proteins are targeted to PicA for translocation across the phage nuclear shell.

### PicA is essential for phage nucleus maturation and phage replication

PhiKZ PicA T367P mutant phage have a delay in importing PhiKZ proteins and consequently a delay in phage nucleus growth. To determine if this delay affects the timing of the PhiKZ lytic cycle, we performed a single step lysis curve in which we monitored cell lysis with a cell impermeable fluorescent DNA dye, SYTOX Green, as a reporter for the completion of the phage lytic cycle (38). Cells infected with wildtype PhiKZ lysed at an average of 62 mpi (n=12) as determined by the time at which 50% maximal fluorescence was reached, whereas lysis of cells infected with PhiKZ PicA T367P was delayed until an average of 75 mpi (n=12) (Fig 4A). Despite this delay in lysis time, the mutant phage displayed no decrease in titer when plated on wildtype *P. aeruginosa* (Fig 4B). The delay in completing the lytic cycle together with the delay in phage nucleus growth suggests that the protein trafficking functions provided by PicA are important for efficient PhiKZ replication and raise the question of whether this highly conserved core gene plays an essential role that is conserved in other phages.

**Figure 4.**
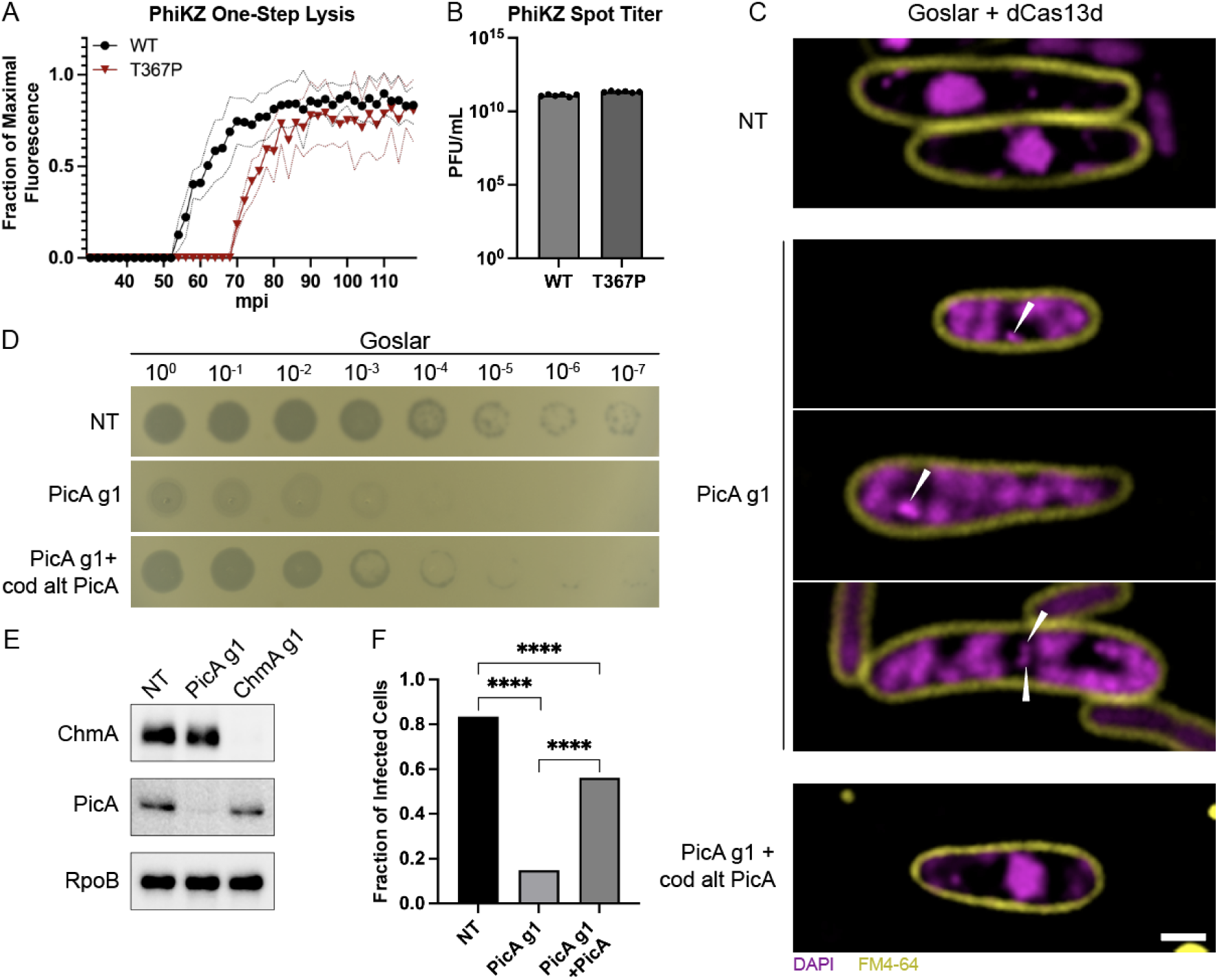
PicA is essential for replication and phage nucleus maturation. A) One-step lysis curves of *P. aeruginosa* infected with wildtype or PicA T367P mutant PhiKZ. Phage induced lysis was detected by Sytox green. Data shown as fraction of maximal fluorescence. Solid line depicts the mean across n = 12 replicates. Dotted lines show mean +/- standard deviation. B) PFU/mL of wild type and PicA T367P mutant PhiKZ as determined by spot titer on lawns of *P. aeruginosa*. N= 5 per condition. C) *E. coli* expressing the indicated constructs (NT = non-targeting) and infected with wildtype Goslar. DNA is visualized with DAPI (purple) and cell membranes are visualized with FM4-64 (yellow). Location of the phage genome in arrested infections is located at the point of the white triangles. Images taken at 75-90 minutes post infection. Scale bar is 1 µm. D) Spot titer of wildtype phage Goslar on lawns of *E. coli* expressing dCas13d with the guides and protein indicated on the left. PicA exogenous expression constructs are codon altered to be insensitive to dCas13d PicA g1 targeting. Dilution of phage from high titer starting lysate (>10^10^ PFU/mL) is indicated above. Experiment was performed in triplicate and a representative image is shown. E) Western blot of Goslar-infected *E. coli* cells expressing dCas13d with the indicated guide RNA at 90 minutes post-infection. Blot was probed sequentially with anti-PicA and anti-ChmA polyclonal antibodies, stripped, and reprobed with anti-RpoB as a loading control. F) Percent of infected cells imaged in (B) that contained a phage nucleus > 0.5 µm (n = 121-182 infected cells per condition). **** = p <.0001 as determined by Fisher’s exact test.

Selection against PhiKZ PicA with catalytically-active Cas13a only produced escape mutants with single nucleotide missense mutations (Table 1), suggesting that the gene encoding PicA is essential. To determine how loss of PicA affects *E. coli* phage Goslar replication, we used the recently developed CRISPRi-ART approach, which utilizes a catalytically-deactivated *Ruminococcus flavefaciens* Cas13d (dCas13d) (39, 40) to selectively knock down Goslar PicA during infection of *E. coli.* We chose to focus on Goslar since we found that the dCas13d repression system was less effective when used in *P. aeruginosa* against PhiKZ.

Spot titers of Goslar on lawns of *E. coli* cells expressing dCas13d and a guide targeting the region surrounding the start codon of PicA (PicA g1) require a ∼10,000-fold higher phage titer for clearing than strains expressing a non-targeting guide (Fig 4D). The clearings in PicA g1 lawns are opaque and do not form individual plaques, making exact efficiency of plating quantification difficult. However the large order of magnitude decrease in phage clearing is readily apparent and consistent across replicates. Complementation of the PicA knockdown with a plasmid-expressed codon-altered *picA* gene that is not targeted by the guide restores phage titer, showing that the effect of the knockdown is specific to PicA (Fig 4D).

We used fluorescence microscopy to understand at which stage infections were blocked in the absence of PicA expression. Individual host cells expressing dCas13d PicA g1 display hallmarks of early infection, including cell swelling and host genome degradation, but are arrested at an early stage of phage replication (Fig 4C). Even by 75-90 mpi, the phage nucleus fails to mature and the phage genome appears as a small DAPI-stained focus in the bacterial cell (Fig 4C, white arrows). Frequently, multiple phage genomes can be seen in cells with arrested infections (Fig 4C). A similar phenotype has previously been observed in infections where the essential nuclear shell protein ChmA is selectively knocked down (40). Western blots of infected cells confirmed that PicA expression was strongly inhibited by PicA g1 expression, but not by the non-targeting guide nor ChmA g1 controls (Figure 4E). Furthermore, ChmA was expressed at approximately wildtype levels during PicA knockdown indicating that the arrested infection is still transcriptionally active (Fig 4E). Complementation of the knockdown with a codon-altered untargetable PicA restores nucleus formation, in agreement with spot titer results (Figs 4C and D), indicating that this phenotype is specific and not due to off-target or polar effects on co-transcribed genes (Fig 4D). Quantification of >100 infected cells per condition shows that a maturing phage nucleus greater than 0.5 micron in diameter forms in the majority (84%) of infected cells expressing the non-targeting guide. However, infected cells expressing the PicA targeting guide only exhibit a maturing phage nucleus 15% of the time. Expressing the non-targetable PicA along with PicA g1 increases the number of maturing phage nuclei present to 56% (Fig 4F). Together, these data indicate that PicA plays an essential role early in nucleus-based phage replication.

## Discussion

### A proposed mechanism for protein trafficking into the phage nucleus

In order to achieve selective protein trafficking into the phage nucleus, the proteins destined for import must be recognized in the cytoplasm and transported across the ChmA shell. We found that the import pathways of PhiPA3 and PhiKZ are highly effective at discriminating between each other’s proteins. We have identified a specific recognition region of an endogenous PhiKZ nuclear protein, gp104, required for transporting it into the nucleus and a conserved shell-associated protein, PicA, that comprises part of the protein import machinery. One of the chimeras generated between PhiKZ gp104 and its homolog PhiPA3 gp108 (PhiPA3gp108-[PhiKZ gp104(113–162)]-sfGFP) revealed a stalled transport intermediate that was targeted to the PhiKZ import machinery, but failed to be transported. The stalled intermediate co-localized with fluorescently tagged PicA foci on the nuclear periphery, suggesting a two-step import model in which proteins are first targeted to PicA and then licensed to be translocated across the nuclear shell by the import machinery (Fig 5A). This process is dependent upon amino acids predicted to be on the surface of the imported protein (Fig 5A), consistent with prior results showing that single amino acid substitutions on the surface of GFPmut1 affect import into the PhiKZ nucleus. When PicA expression is prevented using CRISPRi-ART, phage nucleus development is halted at an early stage of infection, suggesting that the protein import pathway is essential for the phage nucleus-based life cycle.

**Figure 5.**
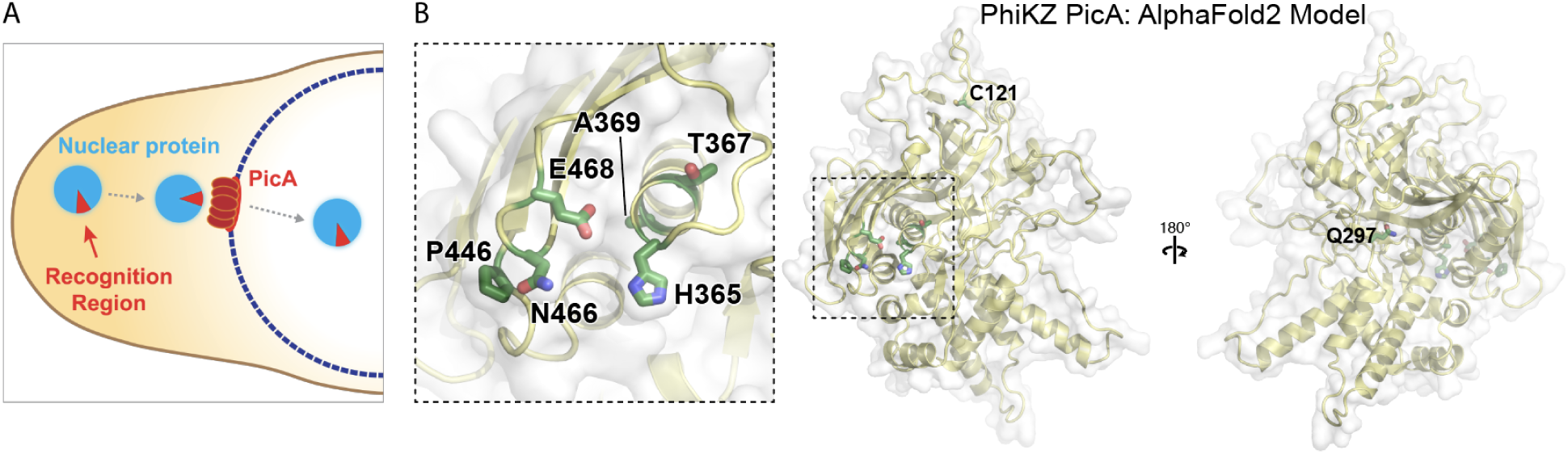
Model for the mechanism of selective protein trafficking into the phage nucleus. A) Mechanistic models for selective protein trafficking into the phage nucleus. B) AlphaFold2 predicted structure of PhiKZ PicA with the location of amino acids which display protein import defects when mutated. There are regions of low confidence for the predicted structure of PicA (see Fig S5), but it is notable that the mutations shown on the left are in the region that is the most confidently predicted.

The PIC pathway is unique compared to other protein translocation systems, such as bacterial TAT and Sec pathways, that depend upon a simple signal sequence at the N- or C-terminus of cargo proteins (24, 27, 28). These cellular systems have evolved to be universal and interchangeable, so one common targeting sequence can be recognized in many divergent species of bacteria, enabling new genes acquired via horizontal gene transfer to be properly localized. In contrast, chimalliviruses face selection pressures from both other chimalliviruses and the host, likely shaping the evolution of distinct import selectivities. The PIC pathway relies on multiple amino acids on the surfaces of both the folded cargo proteins and PicA for import. This provides high substrate selectivity that can be maintained and evolved such that competing phages like PhiPA3 and PhiKZ can coexist in the same virocell without importing each other’s proteins (Fig 1A) (13). We have recently shown that PhiPA3 and PhiKZ frequently coinfect and compete with one another (13, 33). Indeed, importing PhiPA3’s homing endonuclease gp210 into PhiKZ’s nucleus is fatal for PhiKZ (Fig 2) (13, 33). Thus, toxic proteins produced by competing chimalliviruses provide a strong selective pressure to evolve import systems with distinct selectivities (Fig 2). The selectivity of the PIC pathway also protects the phage genomes from host defense proteins, preventing import of DNA-targeting CRISPR-Cas and restriction enzymes (8, 9). Notably, mutations in host defense factors that overcome this protection by targeting to one phage’s nucleus would not defend against other nucleus-forming phages. Thus, the overall mechanism described here requiring two steps (targeting and licensing) and multiple surfaces of cargo and transporter ensures import specificity, protecting these phages from competing viruses and host defenses, and likely plays an important role in shaping the evolution of nucleus-forming phages.

AlphaFold2 (30) produces a predicted structure of PhiKZ PicA with regions of high confidence (Fig 5B, Fig S6). Foldseek and DALI searches find no structural homologs, suggesting this protein forms a novel fold (41, 42). Mapping the mutations generated in this study onto the predicted structure of PicA provides insights into the regions important for selective trafficking (Fig 5B). Six mutations, A369T, P446S, N466K, E468K, H365Q and T367P, all occur within ∼13 angstroms of each other in structural space despite being distant from each other in the primary sequence. Two other mutations, Q297K and C121Y, occur on surfaces distal to each other and the other mutations isolated. (Fig 5B). While the T367P mutation imparts a general defect on protein trafficking, the rest of these mutations provide a deficit in trafficking only one or a small number of nuclear proteins (Fig 5B). Together these mutations demonstrate that residues on distal regions of PicA’s surface play critical and differing roles in cargo selection.

The presence of a pore-like structure in the phage nuclear shell for macromolecular transport has been postulated since the discovery of the phage nucleus (1, 43), but pores large enough to accommodate protein trafficking have yet to be observed in tomograms of infected cells (1, 3–5, 10). One explanation for this is that only a limited number of protein trafficking complexes exist on the nuclear shell. Indeed, sfGFP-tagged PicA forms a single punctum on the nuclear periphery of three different phages, suggesting a higher order oligomeric state. AlphaFold2 does not produce high confidence oligomers of PicA, which likely suggests that other yet-to-be-identified proteins are required for the formation of the protein trafficking complex. The observation of a single PicA punctum that co-localizes with a stalled transport intermediate does suggest though that there may be only a single complex on the nuclear shell where protein trafficking occurs.

## Materials and Methods

### Bacterial strains, phage, and growth conditions

*P. aeruginosa* PAO1 K2733 was used as the host for PhiKZ and PhiPA3. It was grown at 30 ℃ in LB and 15 μg/ml gentamicin was used for plasmid selection. Phage stocks of PhiKZ and PhiPA3 were collected from plate lysates using phage buffer (10 mM Tris (pH 7.5), 10 mM MgSO_4_, 68 mM NaCl, and 1 mM CaCl_2_)) and stored at 4 ℃ with titers of ∼10^11^ PFU/mL. *E. coli* MC1000 was used as the host for Goslar. It was grown at 37 ℃ and 30 μg/ml chloramphenicol and/or 100 μg/ml ampicillin were used for plasmid selection. Phage stocks of Goslar were collected from plate lysates using phage buffer and stored at 4 ℃ with titers of ∼10^10^ PFU/mL.

### Plasmid construction and transformation

Plasmids were synthesized and cloned by Genscript. The vector for all *P. aeruginosa* plasmids was pHERD-30T. The vector for *E. coli* plasmids was either pDSW206 or p15a-CmR. Plasmid transformation was accomplished by 2 kV electroporation of cells washed with 300mM sucrose or 10% glycerol.

### Single cell infection assay

Single cell infections of *P. aeruginosa* and *E. coli* were visualized using fluorescence microscopy. *P. aeruginosa* was grown at 30 °C rotating up to an OD_600_ ∼0.5 and mixed with PhiKZ or PhiPA3 lysate at a ratio of 1:100 lysate:culture. For protein expression, LB was supplemented with 15 μg/ml gentamicin and 0.1% arabinose. Infections were incubated rotating at 30 °C then inoculated on 1% agarose pads containing 25% LB and 1 µg/ml DAPI and dried for 5 minutes at room temperature before a coverslip was applied. Alternatively, *E. coli was* grown at 37 °C to an OD_600_ 0.3 and inoculated on 1% agarose pads containing 25% LB with antibiotics if indicated and grown for 2 hours at 37 °C. For protein expression, agarose pad was supplemented with 0.2 mM IPTG (pDSW206) and/or 25nM aTc (p15A-CmR). 10 μL of Goslar lysate was tadded and incubated at 37 °C. Cells were stained with 25 ug/mL DAPI and 3.75 ug/mL FM4-64 in 25% LB before imaging. Imaging was performed with a DeltaVision Elite Deconvolution microscope (Applied Precision, Issaquah, WA, USA). Images were further processed by the aggressive deconvolution algorithm in DeltaVision SoftWoRx 6.5.2 Image Analysis Program.

### Plaque Assay

For Goslar, 9 mL of 0.35% LB top agar containing antibiotics and inducing agents were combined with 1 mL of overnight *E. coli* culture and poured onto LB plates containing the required antibiotics. Expression of dCas13d was induced with 200nM aTc and expression of PicA was induced with 0.2mM IPTG. A 10-fold dilution series of Goslar was prepared in LB and 2 uL were spotted for each dilution and incubated at 37 °C for 15-18 hours.

For *P. aeruginosa*, liquid cultures were pre-induced with 0.1% arabinose if indicated and grown to an OD_600_ ∼0.5. 200 μL of culture was mixed with 5 mL 0.35% LB top agar +/- 0.1% arabinose and poured onto a plate with the appropriate antibiotic. A 10-fold dilution series of PhiKZ was prepared in phage buffer and 2 uL were spotted for each dilution and incubated at 30 °C for 15-18 hours.

### Efficiency of Plating

Efficiency of plating for mutant PhiKZ was determined by plaque assay on K2733 lawns expressing gp210-GFPmut1 as described above. PFU/mL were determined for wildtype and each mutant. Average titer of wildtype PhiKZ was normalized to one and the efficiency of plating for each mutant was determined relative to wildtype.

### Isolation and sequencing gp210-resistant PhiKZ

100 μl of *P. aeruginosa* K2733 (OD_600_ ∼0.4) induced with 0.1% arabinose was infected with 10 μl of PhiKZ lysate. 5 mL of warm 0.35% LB top agar was added and poured onto an LB plate with 15 μg/ml gentamicin and incubated overnight at 30 °C. Isolated plaques from each plate were then streak purified three more times under the same conditions to obtain clonal phage populations. After the final streak purification, plate lysates were generated in phage buffer. Phage genomic DNA was isolated (see supplement) and whole genome sequencing was performed by SeqCenter (Pittsburgh) using the Illumina NextSeq 2000 platform at a depth of 200Mbp. SeqCenter provided paired end reads (2x151bp) and reported variations from the Genbank entry for PhiKZ (NC_004629.1).

### Isolation of PicA mutant PhiKZ with Cas13a

*P. aeruginosa* K2733 cells containing Cas13a expression vector with the specified guide were grown to OD_600_ ∼0.5 in LB containing 15 μg/mL gentamicin and 1% arabinose. Escape mutants were otherwise isolated as described above. The region of the genome targeted by Cas13a was amplified by PCR and sequenced by Sanger sequencing (Azenta). Whole genome sequencing was performed as described above for a subset of mutants (PhiKZ PicA T367P and Q297K) to determine that no off target mutations were generated.

Alternatively, for PicA guide 6, no plaques could be isolated by the above method without first enriching for mutants in liquid culture. 3 mL of OD_600_ ∼0.5 culture were infected with ∼10^9^ PFU of PhiKZ and grown overnight at 30 °C rotating. Supernatant was collected from the liquid culture the following day and serially diluted until individual plaques could be obtained and purified by the above method.

### Western blot

MC1000 strains were infected on 1% agarose, 25% LB, 6 cm diameter plates containing the appropriate antibiotics and inducers. 75 mpi, cells were collected in 25% LB and ∼4.0 x 10^8^ cells were resuspended in 500 µL 2x SDS loading buffer (0.1 M Tris HCl, 0.004% bromophenol blue, 4% SDS, 20% glycerol, pH 6.8 plus 5% BME) and heated for 4 minutes at 95 °C. 10 µL of each sample were loaded into a Novex 4-20% Tris-Glycine Gel and run at 200V. Protein was transferred to PVDF (Pall Life Sciences) and blocked at room temperature with StartingBlock PBS Blocking Buffer (Thermo Scientific). Membrane was blotted with anti-ChmA (dilution = 1:500; custom polyclonal rabbit antibody generated by GenScript), anti-PicA (dilution = 1:1,000; custom polyclonal rabbit antibody generated by GenScript), and anti-RpoB-HRP (loading control, dilution = 1:5,000; Biolegend) for 1 hour at room temperature with mild agitation. The membranes were washed with TBS-T and incubated with secondary antibody (dilution = 1:10,000; HRP-conjugated goat anti-rabbit IgG (H + L), Invitrogen) (anti-PicA and Anti-ChmA only) for 1 hr at room temperature and washed again. The membranes were visualized via ECL (Cytiva Amersham) using a ChemiDoc MP Imaging System (Bio-Rad). Images were adjusted for figure panels in Adobe Photoshop (21.2.0) and final figures were generated in Adobe Illustrator (24.2).

### Quantification of the fraction of cells with nuclear localization of sfGFP-tagged protein during PhiKZ infections

Single cell infection assay and fluorescence microscopy was performed as described above. The location of the phage genome was determined by locating the approximately circular area of high intensity in the DAPI channel. Nuclear localization of the sfGFP-tagged protein was called if the highest intensity of GFP signal in the GFP channel overlapped with the phage genome in the DAPI channel. Bias was avoided by quantifying every cell imaged across five fields of imaging per condition. N = 68-159 per condition.

### Quantification of Nuclear Cross-Sectional Area

Single cell infection assay and fluorescence microscopy was performed as described above. Nuclear cross-sectional area was determined by tracing the border of the DAPI-stained nuclei in ImageJ at its widest point in the Z-axis. The area was then measured by ImageJ in µm^2^. 60 nuclei were measured per condition, areas were plotted with PRISM, and statistical significance was determined by Mann-Whitney test.

### Quantification of Nuclear Diameter

Single cell infection assay and fluorescence microscopy was performed as described above. Infected cells were identified by the presence of cell swelling and host genome degradation. DAPI-stained foci indicative of phage genomes were measured at their widest point using DeltaVision SoftWoRx 6.5.2 Image Analysis Program. Infected cells were then binned into two categories, those containing a DAPI stained focus greater or less than 0.5 µm. All cells imaged per each condition were quantified to avoid bias. N ranged from 121 to 183 infected cells per condition. Statistical significance was determined by Fisher’s exact test performed as a pairwise comparison between each of the three conditions.

### Single Step Time-to-Lysis

Experiment was performed essentially as described in Egido et Al. (38). In short, *P. aeruginosa* K2733 was grown at 37 °C to OD_600_ ∼0.25 in LB. 30μL of phage lysate (wildtype or mutant PhiKZ) was added to 3mL of bacteria (MOI = ∼5) and incubated at 30°C for 25 minutes. After 25 minutes, Sytox Green Nucleic Acid Stain (ThermoFisher) was added to a final concentration of 5μM. 200 μL of each culture was added to a well of a black-walled, clear-bottom 96-well plate (Costar), with 12 wells per condition. Fluorescence measurements were performed in a microplate reader (Tecan Infinite M Plex) at 30 °C. Fluorescence measurements were taken at: λ_excitation_ = 504 nm; λ_emission_ = 537 nm; gain = 25; flashes per well = 5. Measurements began at 30 mpi and were taken every 2 minutes up to 120 minutes post-infection. Data analysis was done with Microsoft Excel and Prism (GraphPad). For each replicate, background fluorescence was subtracted and values were divided by the maximum fluorescence detected. Negative values due to photobleaching of the background prior to lysis were set to zero. The mean +/- the standard deviation across 12 replicates was plotted and average time-to-lysis determined as the time point at which the mean fraction of maximal fluorescence reached 0.5.

### Fluorescence Colocalization Analysis

We used images from fluorescence microscopy experiments to perform object-based colocalization analyses of PhiPA3gp108[PhiKZgp104(113–162)]-sfGFP and PicA-mCherry foci in PhiKZ infected cells in eight different fields of view (FOV). Two FOVs were excluded from the analysis due to sample drift. The analysis and corresponding scripts were performed and written in MATLAB (see supplement). A total of 187 infected cells had fluorescence above threshold and were included in the analysis.

## Acknowledgements

This work was supported by an Emerging Pathogens Initiative grant from the Howard Hughes Medical Institute (to J.P., K.P, E.V., and K.D.C.), the National Institutes of Health R01-GM129245 (to J.P., E.V., and K.P.), and R35 GM144121 (to K.D.C.). C.J.M. was supported by a National Institutes of Health T32 training grant through the UCSD Medical Scientist Training Program. E. G. A and A. P. were supported by the Pathways in Biological Sciences National Institutes of Health T32 training grant (T32 grant GM133351). W.W. was supported by Second Century Fund (C2F) of Chulalongkorn University. B.A.A. was supported by m-CAFEs Microbial Community Analysis & Functional Evaluation in Soils, a Science Focus Area led by Lawrence Berkeley National Laboratory based upon work supported by the US Department of Energy, Office of Science, Office of Biological & Environmental Research [DE-AC02-05CH11231]. B.F.C. was supported by U. S. Department of Energy, Office of Science, through the Genomic Science Program, Office of Biological and Environmental Research, under the Secure Biosystems Design Initiative project Intrinsic Control for Genome and Transcriptome Editing in Communities (InCoGenTEC).

---

**Supplementary Figure 1.**
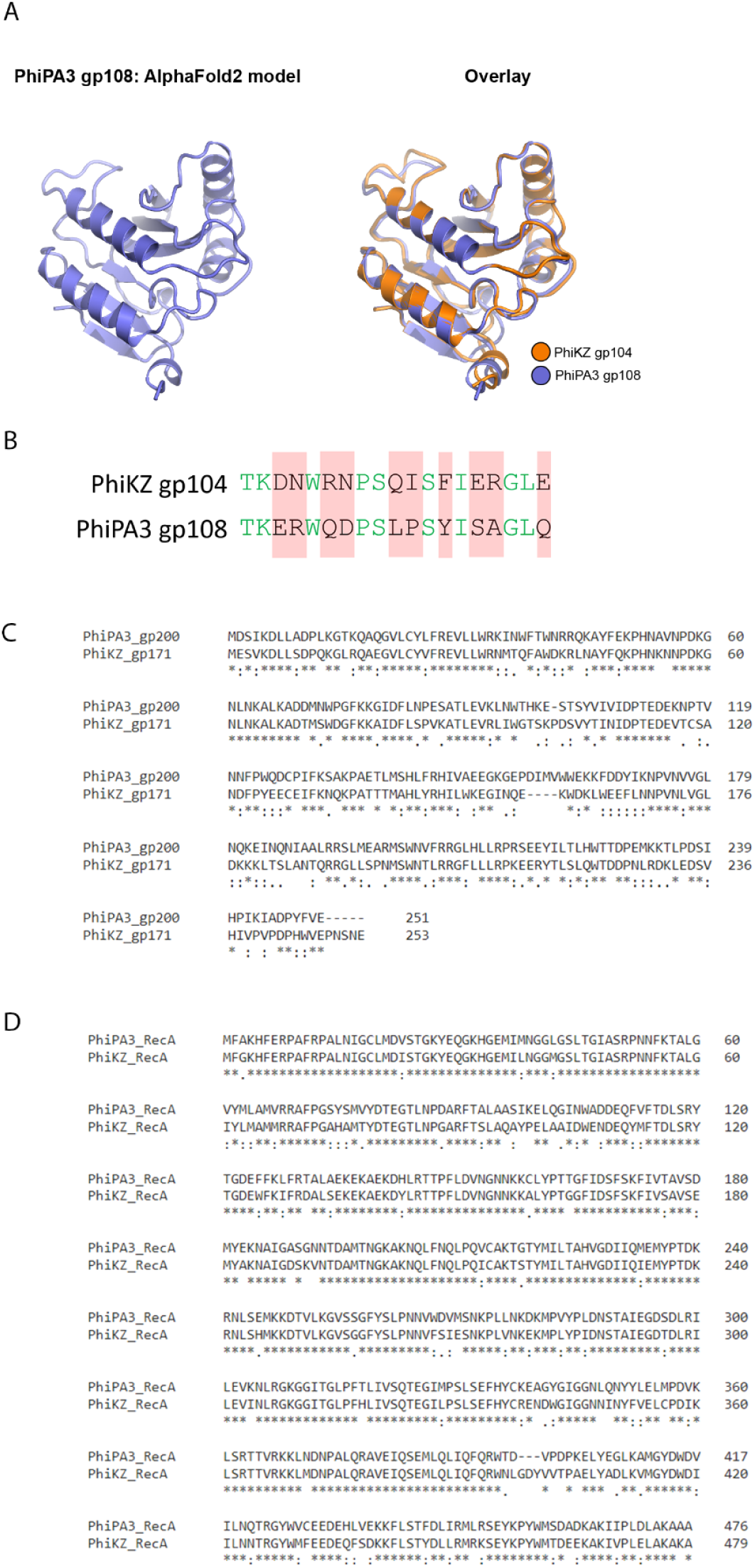
Structural and sequence alignments of PhiKZ and PhiPA3 nuclear proteins. A) AlphaFold predicted structures of PhiPA3 gp108 (left) and structures of PhiKZ gp104 and PhiPA3 gp108 aligned with PyMol (right). RMSD = 0.487 Angstrom. B) PhiKZ gp104 amino acids 77-95 aligned with the corresponding region in PhiPA3 gp108. Conserved amino acids are in green. Non-conserved amino acids are in black with red highlighting. C) Clustal Omega generated sequence alignment of PhiPA3 gp200 and PhiKZ gp171. D) Clustal Omega generated sequence alignment of PhiPA3 RecA and PhiKZ RecA.

**Supplementary Figure 2.**
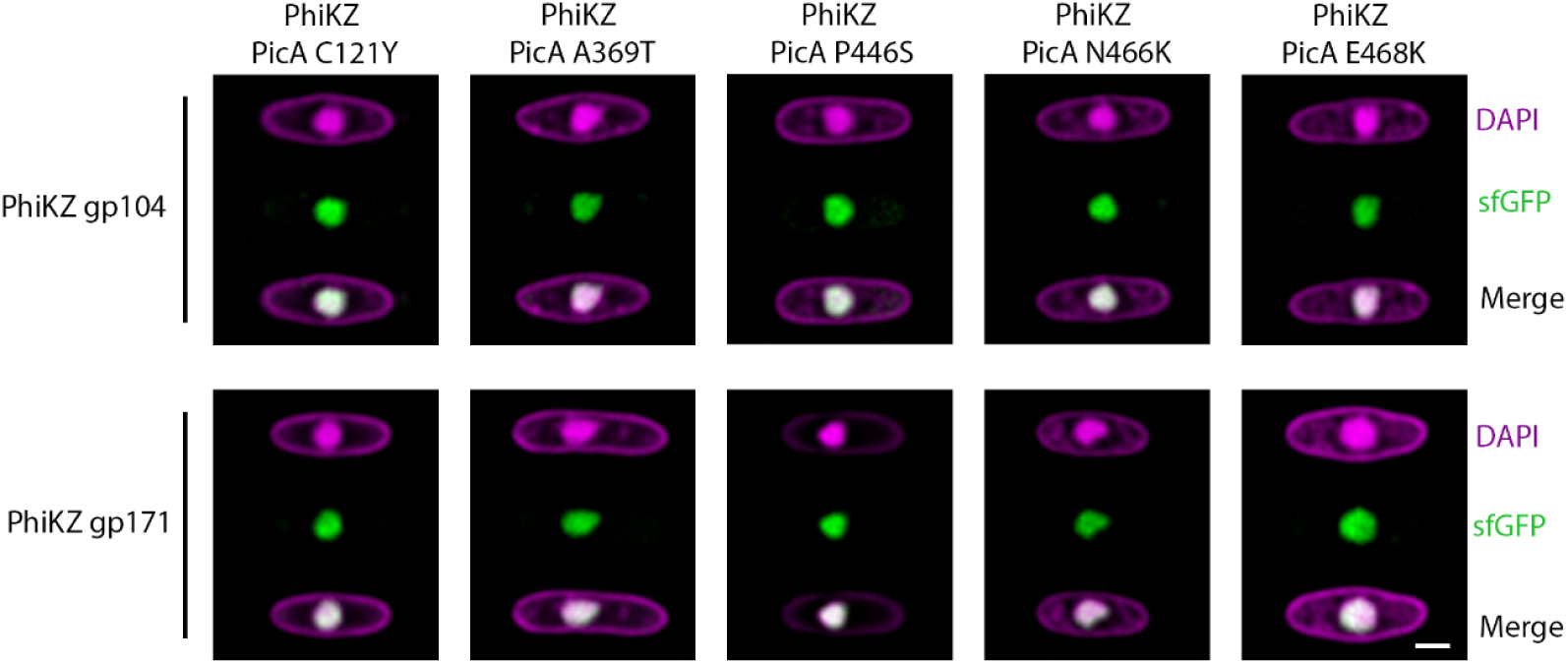
PicA mutant PhiKZ that exclude GFPmut1 still import phage proteins. PicA mutant PhiKZ infecting *P. aeruginosa* cells expressing sfGFP-tagged PhiKZ gp104 (upper panels) or PhiKZ gp171 (lower panels). Scale bar is 1 µm.

**Supplementary Figure 3.**
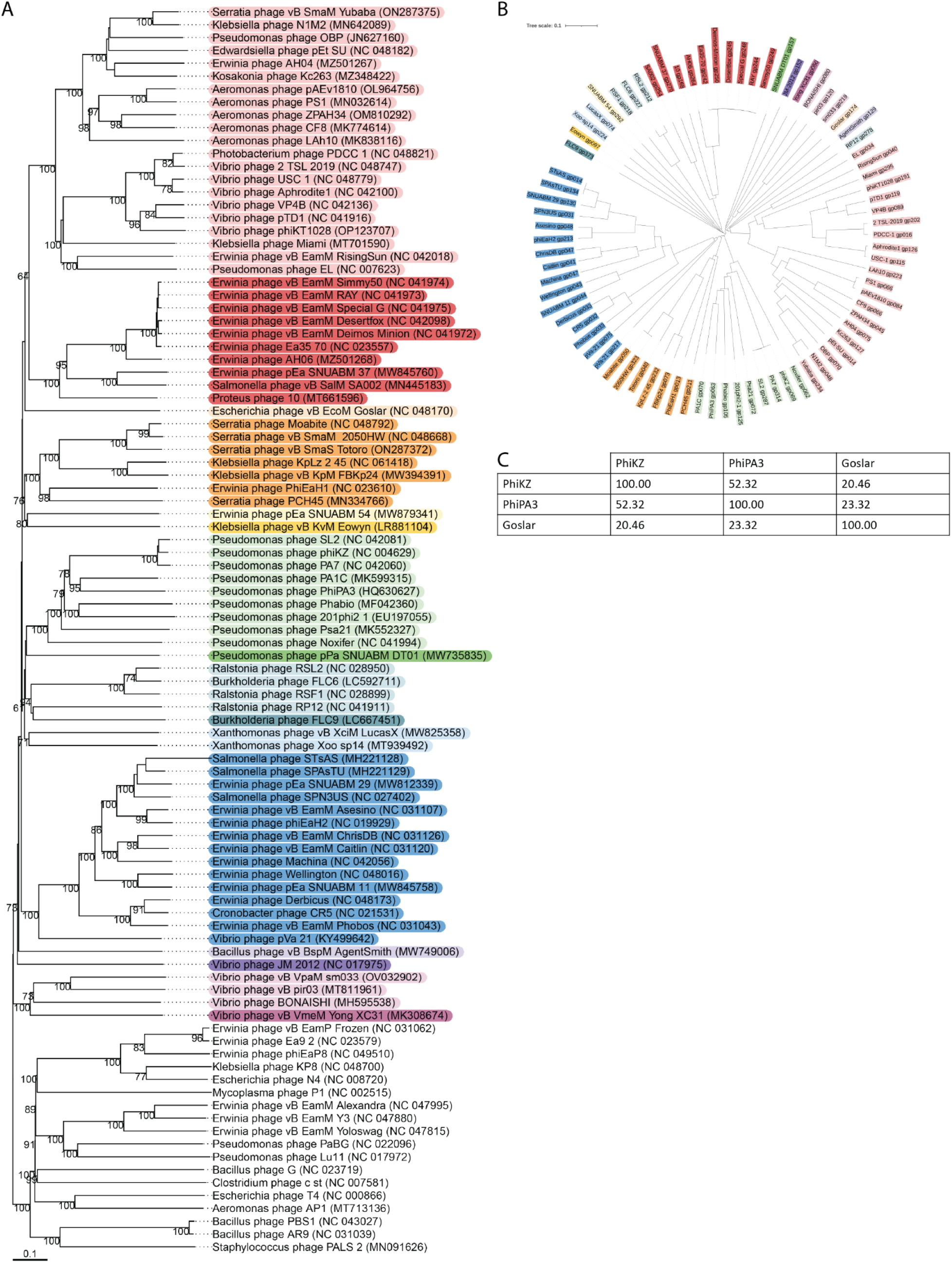
Phylogenetic Analysis of PicA. (A) A phylogenetic tree comparing the whole genomes of all chimalliviruses, showing that they form a single clade. As PicA is part of the core genome of chimalliviruses, homologs are encoded in all members of this group. Chimalliviruses are colored in rainbow by VICTOR-predicted genus groups. (B) A phylogenetic tree of PicA homologs. In general, the protein tree is congruent with the genome tree with the two exceptions of *Erwinia* phage SNUABM_54 clustering with *Xanthomonas* phages Xoo-sp14 and LucasX rather than by itself as a singleton and *Ralstonia* phage RP12 being separate from *Ralstonia* phages RSL2 and RSF1 and *Burkholderia* phage FLC6. Interestingly, *Vibrio* phage pVa-21 has two homologs of this protein, one with a long tail and one without. (C) A comparison of the % identity of the amino acid sequence of the PicA homologs from PhiKZ (gp69), PhiPA3 (gp63), and Goslar (gp174).

**Supplementary Figure 4.**
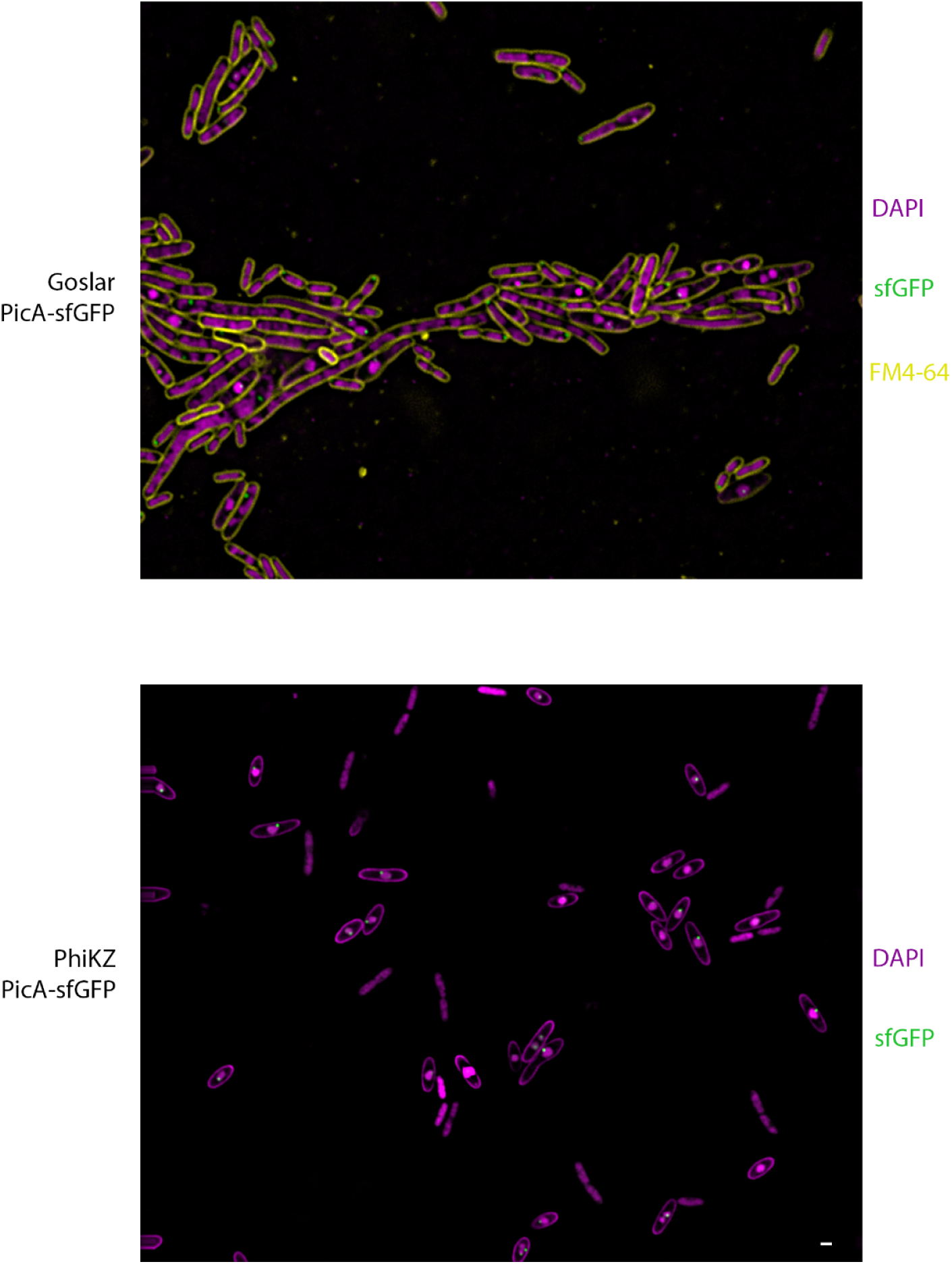
Cells expressing PicA-sfGFP infected with phage. *E. coli* cells expressing PicA-sfGFP infected with Goslar (upper panel) or *P. aeruginosa* cells expressing PicA-sfGFP infected with PhiKZ (lower panel). Scale bar is 1 µm.

**Supplementary Figure 5.**
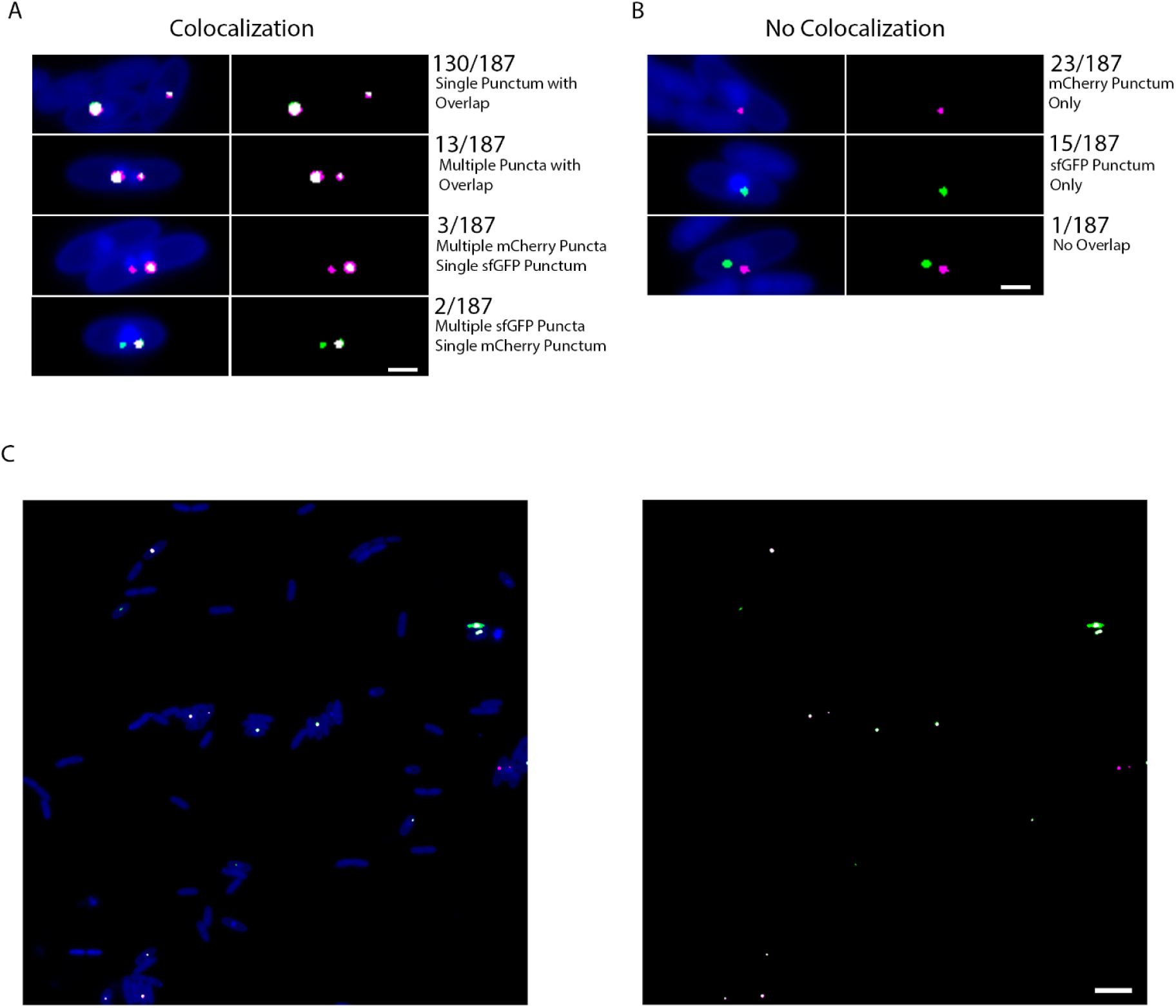
Colocalization Analysis of PhiPA3gp108-(PhiKZ gp104 113-162)-sfGFP and PicA-mCherry. A) *P. aeruginosa* cells co-expressing the chimera PhiPA3gp108-(PhiKZ gp104 113-162)-sfGFP and PicA-mCherry infected with PhiKZ imaged at 30-45 minutes post infection. Images were analyzed by a non-biased object-based colocalization algorithm and then grouped into the colocalization phenotypes shown. The total number of cells with that phenotype is listed on the right. DAPI is shown in blue, sfGFP signal above threshold is shown in green, mCherry above threshold is shown in magenta. Pixels that contain both mCherry and sfGFP above threshold are white. Total number of infected cells with fluorescence above threshold is 187. Scale bar is 1 µm. B) Same as described in (A) except phenotypes shown for cells that did not have overlapping fluorescence are shown. Scale bar is 1 µm. C) Representative whole microscopy field. A total of 8 microscopy fields were analyzed. 2 microscopy fields were left out of the analysis due to drift between sampling of fluorescence channels. Scale bar is 5 µm.

**Supplementary Figure 6.**
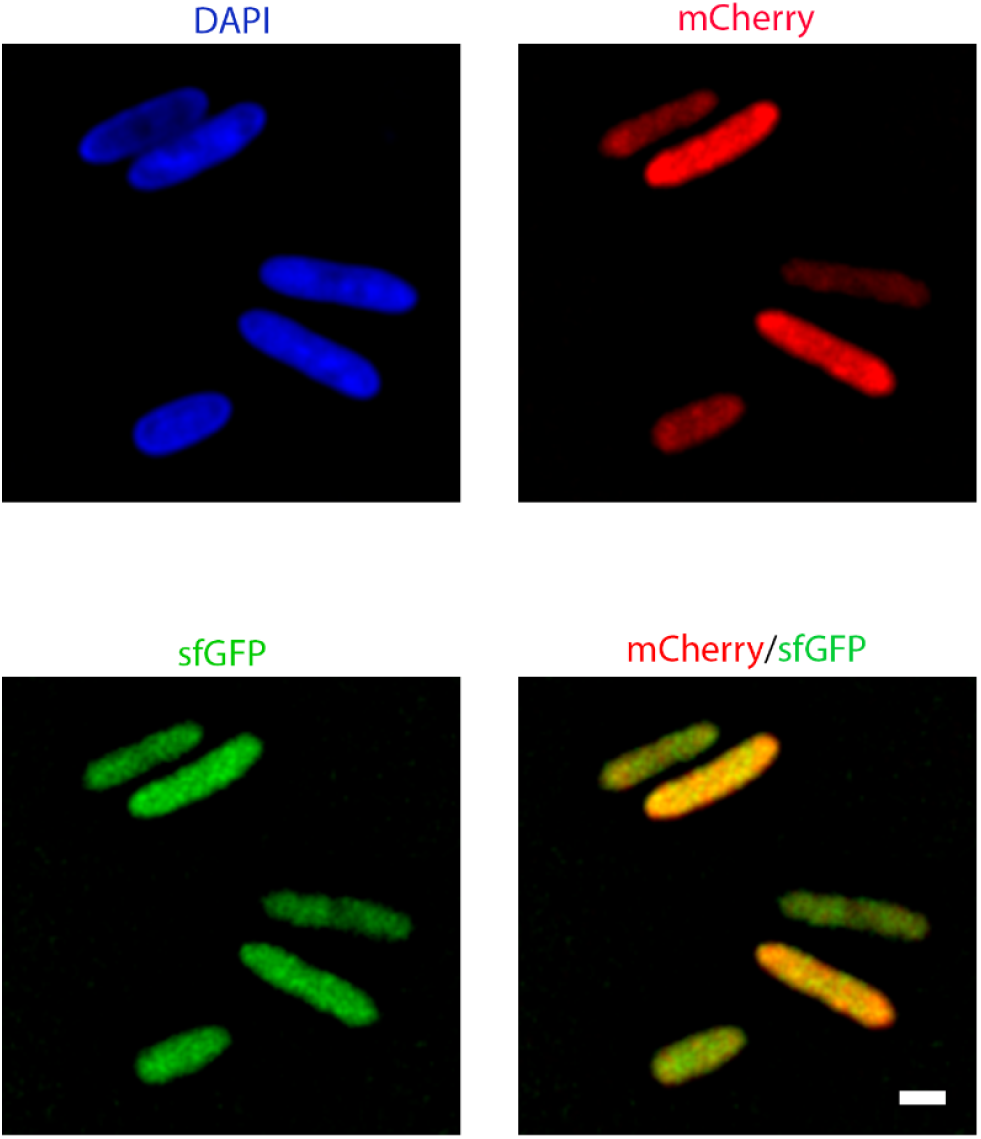
PicA-mCherry and PhiPA3gp108-[PhiKZ gp104(113–162)]-sfGFP are diffuse in uninfected cells. Microscopy image of *P. aeruginosa* cells expressing both PicA-mCherry and PhiPA3gp108-[PhiKZ gp104(113–162)]-sfGFP. Scale bar is 1 µm.

**Supplementary Figure 7.**
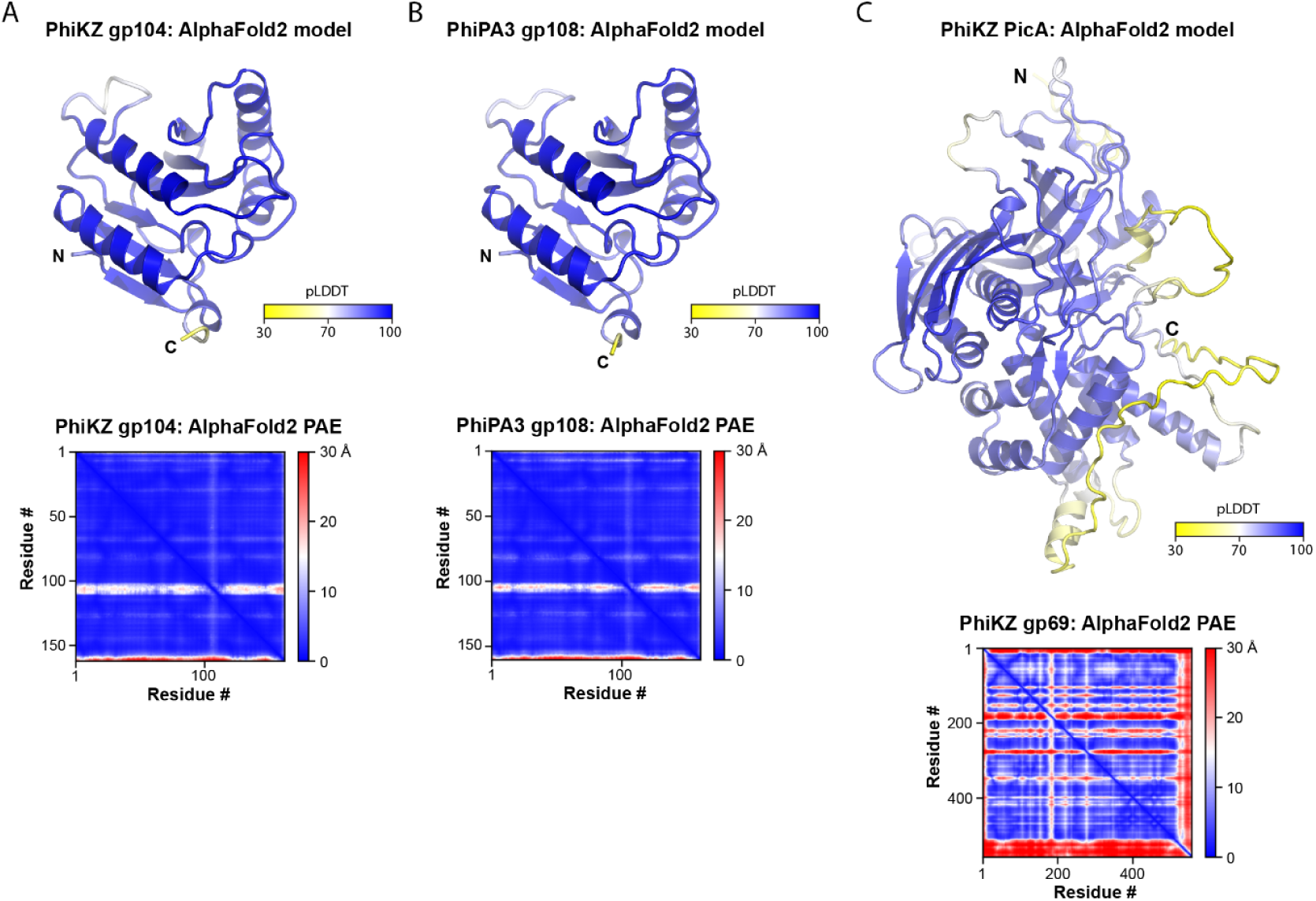
AlphaFold2 predicted structures and predicted aligned errors. Alphafold2 predicted structure colored by pLDDT and predicted aligned error chart for PhiKZ gp104 (A), PhiPA3 gp108 (B) and PhiKZ PicA (C).

**Supplementary Table 1.**
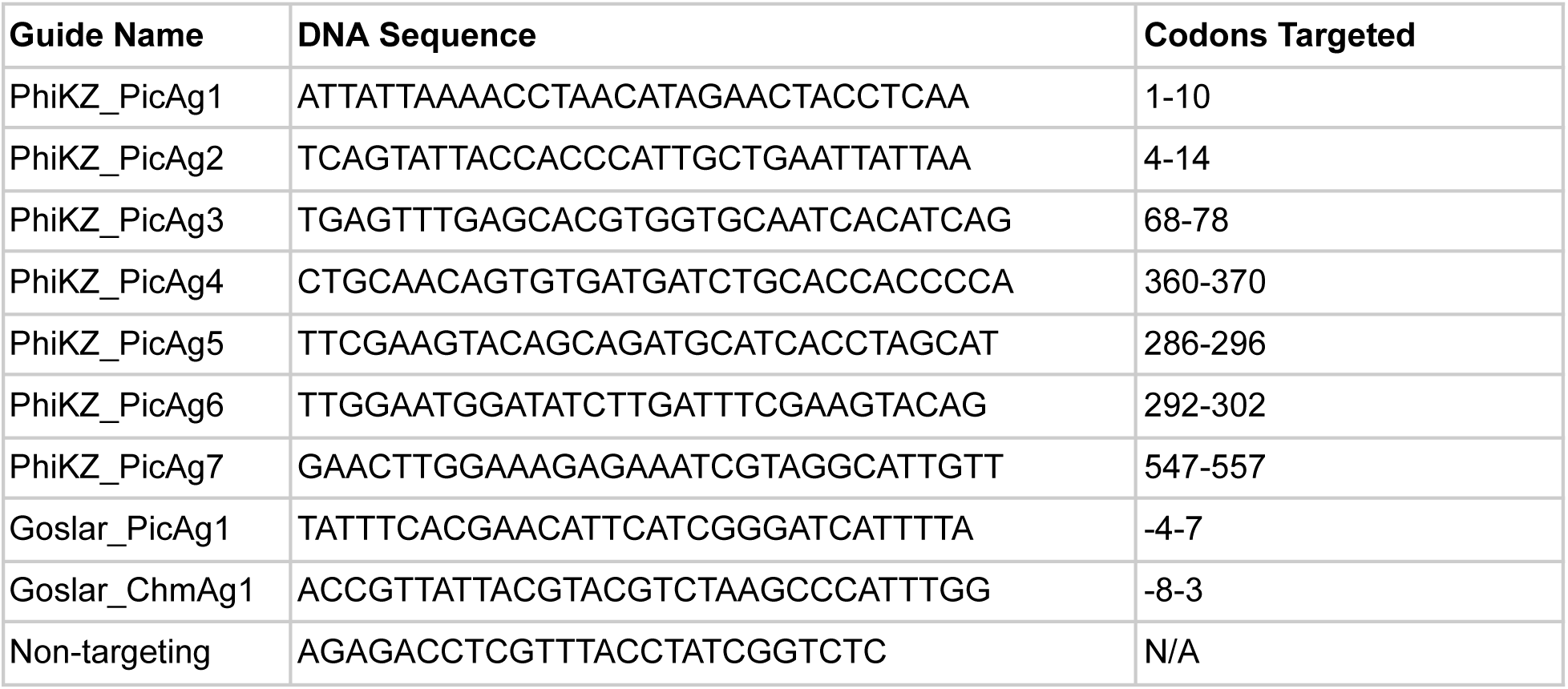
Cas13 guide RNAs used in this study.

**Supplementary Table 2.** Plasmid constructs used in this study.

| **Backbone** | **Insert** |
| --- | --- |
| pHERD30T | PhiPA3_RecA-sfGFP |
|  | PhiPA3_gp108-sfGFP |
|  | PhiPA3_gp200-sfGFP |
|  | PhiPA3_gp78-sfGFP |
|  | PhiPA3_gp257-sfGFP |
|  | PhiKZ_RecA-sfGFP |
|  | PhiKZ_gp104-sfGFP |
|  | PhiKZ_gp171-sfGFP |
|  | PhiKZ_gp104(1-113)-sfGFP |
|  | PhiKZ_gp104(66-113)-sfGFP |
|  | PhiKZ_gp104(66-162)-sfGFP |
|  | PhiPA3_gp108[PhiKZ_gp104(1-65)]-sfGFP |
|  | PhiPA3_gp108[PhiKZ_gp104(66-162)]-sfGFP |
|  | PhiPA3_gp108[PhiKZ_gp104(66-115)]-sfGFP |
|  | PhiPA3_gp108[PhiKZ_gp104(66-95)]-sfGFP |
|  | PhiPA3_gp108[PhiKZ_gp104(77-100)]-sfGFP |
|  | PhiPA3_gp108[PhiKZ_gp104(66-76)]-sfGFP |
|  | PhiPA3_gp108[PhiKZ_gp104(79-88)]-sfGFP |
|  | PhiPA3_gp108[PhiKZ_gp104(96-115)]-sfGFP |
|  | PhiPA3_gp210-GFPmut1 |
|  | PhiPA3_gp210-gp104 |
|  | GFPmut1 |
|  | PhiKZ_PicA + GFPmut1 |
|  | PhiKZ_PicA-sfGFP |
|  | PhiPA3_PicA-sfGFP |
|  | Lbu_Cas13a_PicAg1 |
|  | Lbu_Cas13a_PicAg2 |
|  | Lbu_Cas13a_PicAg3 |
|  | Lbu_Cas13a_PicAg4 |
|  | Lbu_Cas13a_PicAg5 |
|  | Lbu_Cas13a_PicAg6 |
|  | Lbu_Cas13a_PicAg7 |
|  | PhiKZ_PicA-mCherry + PhiPA3gp108[PhiKZgp104(113-162)]-sfGFP |
| pDSW206 | Goslar_PicA-sfGFP_codonaltered(1-30) |
|  | Goslar_PicA_codonaltered(1-30) |
| p15A-CmR | dCas13d_Goslar_ChmAg1 |
|  | dCas13d_Goslar_PicAg1 |
|  | dCas13d_Non-targeting |

## Supplementary Methods

### Phage Genomic DNA Isolation

Phage genomic DNA was isolated using 10 mL of lysate first incubated with 5 μl of each RNaseA (100 mg/mL) and DNaseI (20mg/mL) at 37°C for 30 minutes and then 4 ml of phage precipitant (30% PEG 8000, 19.3% NaCl in ddH_2_O) was added and incubated overnight at 4 °C. Samples were centrifuged at 10,000 rcf 4°C for 20 minutes and resuspended in 0.5 ml sterile water. 2.5 ml of Qiagen Buffer PB was added and incubated at RT for 10 minutes. The resuspensions were then filtered through Qiagen PCR Purification columns as specified by the manufacturer and then eluted with 100 uL of sterile water.

### Fluorescence Colocalization Analysis

https://github.com/koeinlow/Colocalization_analysis_Pogliano_lab/. In general, pairs of images corresponding to a specific FOV showing sfGFP and mCherry fluorescence were globally thresholded and binarized. For 2-D pixel-based object detection, we identify connected components (objects) using 4-connected pixel neighborhoods. Objects which contain less than 5 and more than 150 pixels are removed, and corresponding label matrices are generated for each discrete object. Image pairs are overlaid to create a composite image, and label matrices are again generated for overlapping objects and pixels. Composite images were then overlaid onto DAPI-stained cells to determine the number of infected cells that display overlapping or distinct fluorescence in one or both channels. Cells were then binned into the phenotypes shown in Figure S5 and the number of cells displaying each phenotype were counted.

### AlphaFold Structural Prediction

Structural predictions were generated with AlphaFold2 (30) software accessed through Google Colab notebook (https://colab.research.google.com/github/sokrypton/ColabFold/blob/main/AlphaFold2.ipynb). Amino acid sequences used for AlphaFold2 structural predictions were retrieved from the National Center for Biotechnology Information database. Structures were visualized and figures generated with PyMOL software (The PyMOL Molecular Graphics System, Version 2.0 Schrödinger, LLC).

### Phylogenetic Analysis

All phylogenetic analyses performed compare protein sequences. The genome-wide phylogenetic tree was made using VICTOR (44), and the trees made for individual proteins were made using Clustal Omega 2.1 (31) and annotated in iTOL (45). Percent identity values were calculated from Clustal Omega alignments.

